# Regional differences in the abiotic environment contribute to genomic divergence within a wild tomato species

**DOI:** 10.1101/744797

**Authors:** Matthew JS Gibson, Leonie C Moyle

## Abstract

The wild currant tomato *Solanum pimpinellifolium* inhabits a wide range of abiotic habitats across its native range of Ecuador and Peru. Although it has served as a key genetic resource for the improvement of domestic cultivars, little is known about the genetic basis of traits underlying local adaptation in this species, nor what abiotic variables are most important for driving differentiation. Here we use redundancy analysis (RDA) and other multivariate statistical methods (structural equation modeling (SEM) and generalized dissimilarity modeling (GDM)) to quantify the relationship of genomic variation (6,830 single nucleotide polymorphisms) with climate and geography, among 140 wild accessions. RDA, SEM, and GDM each identified environment as explaining more genomic variation than geography, suggesting that local adaptation to heterogeneous abiotic habitats may be an important source of genetic diversity in this species. Environmental factors describing temporal variation in precipitation and evaporative demand explained the most SNP variation among accessions, indicating that these forces may represent key selective agents. Lastly, by studying how SNP-environment associations vary throughout the genome (44,064 SNPs), we mapped the location and investigated the functions of loci putatively contributing to climatic adaptations. Together our findings indicate an important role for selection imposed by the abiotic environment in driving genomic differentiation between populations.

## INTRODUCTION

Local adaptation is thought to underlie a large proportion of phenotypic variation segregating within species. Despite this, the primary environmental agents driving selection between natural populations are still unknown in most systems. Across landscapes, genetic divergence can result from selection imposed by environmental forces, but also from the effects of restricted gene flow and genetic drift when subpopulations are partially isolated (Wright, 1943). Disentangling the concurrent effects of spatial isolation and natural selection on generating genetic diversity is critical for understanding the relative importance of adaptation in driving diversification, as well as for identifying the environmental factors underlying its occurrence. This remains a major technical challenge, mainly because collinearity is common among spatial and environmental variables (Kissoudis et al., 2016; Lasky et al., 2012; Lee & Mitchell-Olds, 2011; Wiens, 1989). New statistical methods and the increasing availability of fine-scale environmental and genomic data provide one approach to estimate the independent contributions of environment and space to explaining patterns of genetic variation. These strategies can complement traditional tests for local adaptation (i.e., reciprocal transplants) and begin to characterize the nature of selection acting in the wild, as well as its genetic targets (Fournier-Level et al., 2011; Hancock et al., 2011; Lasky et al., 2012, 2015).

*Solanum pimpinellifolium* L. (currant tomato) is a diploid, short-lived perennial relative of domesticated tomato (*Solanum lycopersicum* L.) that has historically served as a key resource for the identification and breeding of agronomically-important traits (Pedley & Martin, 2003; Pitblado & Kerr, 1979; Tanksley, 2004; Tanksley et al., 1996). Population germplasm collections (accessions) are diverse, varying substantially in mating system (higher rates of outcrossing in the center of the range; Rick et al., 1977) as well as in tolerance to acute abiotic stressors such as salinity and drought (Bolarin et al., 1991; Cuartero et al., 1992; Kissoudis et al., 2016; Foolad, 2007; Foolad & Yin, 1997; Razali et al., 2018). Previous studies have suggested that a large fraction of this phenotypic variation could be attributable to spatially varying selection imposed by the abiotic environment (e.g., Zuriaga et al., 2009; Blanca et al., 2012, 2015). *S. pimpinellifolium* occupies a large geographic range from Ecuador to southern Peru (Figure 1); populations therefore experience environments that vary from cool mid-elevation habitats in the Andes to coastal deserts and tropical rainforests along the pacific ocean, and the associated intraspecific variation in abiotic conditions(climate and soil characteristics; Figure 1) is proposed to shape numerous ecologically relevant traits (Chitwood et al., 2012a, 2012b; Nakazato et al., 2008; Nakazato et al., 2010; Rick et al., 1977; Zuriaga et al., 2009). For example, using common garden experiments Nakazato et al. (2008) identified significant intraspecific variation for drought tolerance between accessions from coastal and highland regions, consistent with a past history of selection that ranges from water restricted (and drought tolerant) on the coast to less restricted (and less tolerant) in the highlands. More broadly, using Mantel tests Blanca et al. (2015) showed that major patterns of genetic structure among *S. pimpinellifolium* accessions correspond strongly with regional bioclimatic variation, and a clear pattern of latitudinal population structure has been demonstrated in this species several times (Blanca et al. 2012, 2015; Lin et al., 2019; Rick et al., 1977; Zuriaga et al., 2009). Between wild tomato species, distribution models have further suggested that local climatic variation (especially in temperature and water availability), rather than geographic isolation, is the main factor explaining contemporary distributions (Nakazato et al., 2010). However, while these studies point to the potential importance of abiotic selection in shaping patterns of intraspecific genetic variation, the degree to which population genetic structure is determined by environmental factors rather than demographic history, the key selective variables responsible, and the identity of relevant/causal loci remain unknown. Describing and understanding these factors is essential for uncovering the processes that initiate phenotypic divergence, including the relative role of local adaptation in driving diversification and the number and identity of its genetic targets.

**FIGURE 1.**
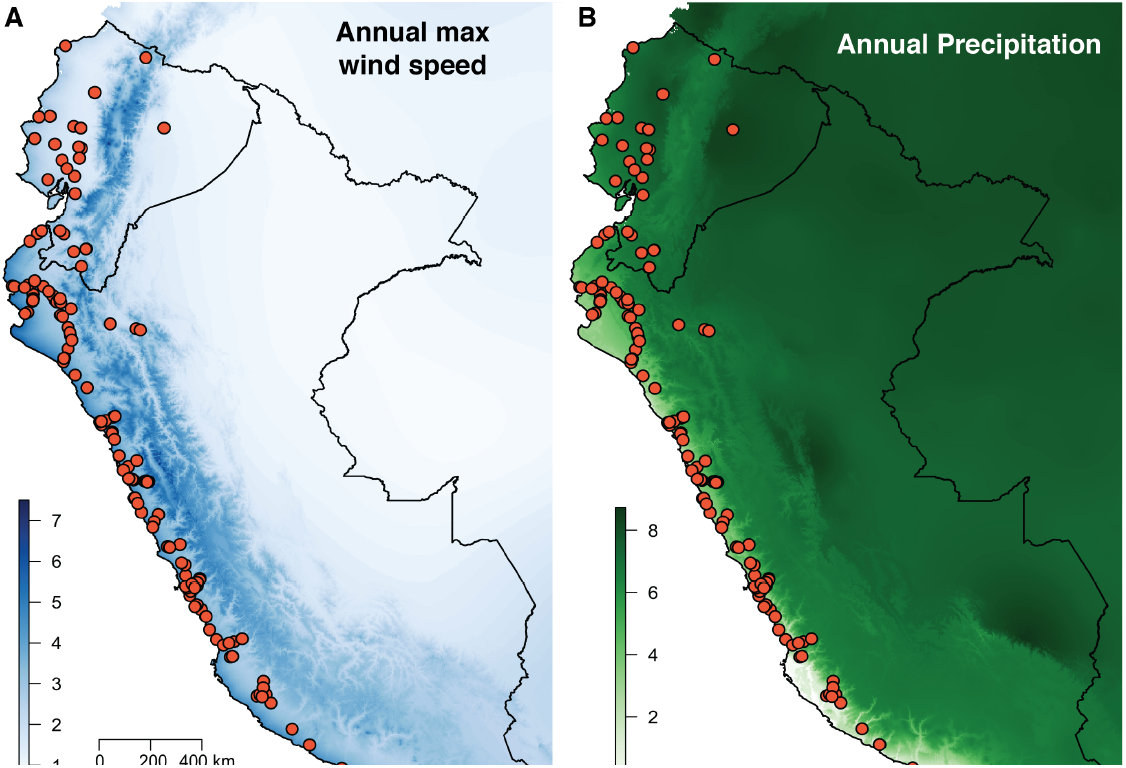
140 South American accessions included in our analyses. Ecuador (North) and Peru (South) are outlined in black. (A) Accession collection sites overlain on average vapor pressure (in units of kPa). Variation in vapor pressure contributed the most to explaining patterns of spatial genetic variation using RDA. (B) Accessions overlain on precipitation seasonality (coefficient of monthly variation).

The main goals of this study were to quantitatively characterize the environmental forces that might impose selection on ecologically relevant traits across the range of *S. pimpinellifolium*, to determine the degree to which these environmental factors explain genetic variation independent of demographic history, and to identify potential genetic targets (SNPs and genes) of this environmental selection. In the absence of confounding population structure, associations between allele frequencies and environmental variables are considered evidence for local adaptation (Endler, 1986). These associations can be used to identify both the forces responsible for imposing selection, as well as the specific loci involved in adaptive evolutionary responses. However, since both adaptation and spatially structured gene flow can result in correlations between allele frequencies and the environment (Endler, 1977), demographic history can cloud such inferences of local adaptation. Moreover, abiotic selection is often the result of a combination of many partially colinear variables, making simple gene-environment correlations incomplete and potentially misleading. To address these issues, multivariate statistical methods commonly used in community ecology—in particular the multivariate regression technique “redundancy analysis” (RDA; Legendre & Legendre, 1998) and other constrained ordination procedures—have recently been applied to landscape genomics to study how climatic variation affects spatial patterns of genomic variation (Lasky et al., 2012, 2016; Forester et al., 2018; Wang et al., 2013). These methods have been used to identify both the sources of selection imposed by the multivariate environment and their genetic targets (individual loci) (Forrester et al., 2018; Lasky et al., 2012, 2016; Lee & Mitchell-Olds, 2011; Manel et al., 2010; Salathe & Schmid-Hempel, 2011; Sork et al., 2010). In doing so, they have begun to address the challenges of disentangling geographical structure from environmental variation in order to quantify the independent contribution of environmental forces to shaping genomic variation across space.

Here we employ RDA to study the relationship between environmental and SNP data matrices while modeling population structure as a covariate, in order to estimate the relative influence of abiotic adaptation on genomic variation across the native range of *S. pimpinellifolium*. Specifically, we use RDA to 1) estimate the independent quantitative contributions of climate, space, and their colinear portion to explaining patterns of genome-wide SNP variation; 2) characterize the specific abiotic variables with the largest influence on genetic variation; and 3) identify genomic regions showing strong associations with major axes of multivariate climate and investigate their potential functional roles in mediating local adaptation. We complement these RDA analyses with other methods designed to quantify the roles of selection and spatial isolation. Our analyses reveal that models including several key abiotic variables explain more genomic variation among genotypes than models considering only geographic isolation. They also allow us to identify loci that have significant associations with environmental variation, after incorporating population structure via partial regression. Our results suggest a prominent role for local adaptation in driving genetic divergence among populations of this agronomically important crop wild relative, and provide a substantial step towards dissecting the contributions of specific loci to local adaptation.

## 2 METHODS

### 2.1 Plant material and genotyping

Seeds from 140 wild georeferenced *S. pimpinellifolium* accessions (Figure 1; Table S1) spanning the entire species range were obtained from the Charles M. Rick Tomato Genetic Resource Center (TGRC; Davis, Calif.). Seeds were soaked in a 50% bleach solution (as per the TGRC protocol), germinated, and transplanted into 1-gallon pots at the Indiana University greenhouses. Fresh leaf material was collected from one plant per accession at 30 days post-germination. DNA was extracted using DNeasy Plant Mini Kits (Qiagen, Valencia, Calif, USA) and two double-digest restriction site associated sequencing (ddRAD) libraries were prepared using *PstI* and *EcoRI* enzymes by the Indiana University Center for Genomics and Bioinformatics. These enzymes were selected to enrich for cut sites in coding regions based on an in silico digestion of the *S. lycopersicum* reference genome ver. SL3.0 (Tomato Genome Consortium, 2012). Libraries were sequenced across two Illumina NextSeq flowcells (150 bp, paired-end, high-output). Raw reads were trimmed of Illumina adapter sequences using *fastp* (adapter type was automatically detected with fastp; Chen et al., 2018), and then demultiplexed by individual using the process_radtags program in Stacks (ver. 2; Catchen et al., 2013). Reads were processed in sliding windows (15% of read length). Windows with scores below 5 and reads less than 30 bp were discarded. Sequencing error was high at restriction sites due to low base diversity and were changed to the correct restriction overhang (*PstI*; 5‘-TGCA-3‘) to avoid mapping errors. fastp was also used to correct low quality base calls in overlapping regions of paired-end reads (minimum of 30bp overlap required for correction). Read quality was verified pre- and post-processing using fastqc Andrews, 2010). After demultiplexing, reads were mapped to the *S. lycopersicum* reference genome version SL3.0 using default parameters in BWA (Li & Durbin, 2009). Only uniquely mapped paired-end reads were retained (filtering performed with samtools; Li et al., 2009). Variants were called using the Stacks ref_map pipeline, using alpha of 0.05 to call variant sites and genotypes. Genotype calls made with fewer than 4 reads were changed to missing with vcftools and only bi-allelic SNPs typed in at least 30 percent of individuals (42/140) were retained. Further filtering was applied prior to running specific analyses (see below). All pipeline scripts are available at http://github.com/gibsonmatt/pimpGEA.

### 2.2 Environmental data

For each accession, data were compiled for 54 abiotic environmental parameters available from public databases. 19 bioclimatic variables as well as monthly solar radiation, water vapor pressure, and wind speed data were obtained directly from the WorldClim2 database (Fick & Hijmans, 2017). Additional transformations of the WorldClim data were obtained from the Climate South America (ClimateSA v1.0; Hamann et al., 2013) database. Global aridity measures were obtained from the Consultative Group on International Agricultural Research (CGIAR; Zomer et al., 2008). Lastly, soil pH, type, density (fine earth fraction), and water capacity at a depth of 5cm were obtained from the SoilGrids initiative (Hengl et al., 2017). All variables were centered, scaled, and tested for normality using Shapiro-Wilks tests. If appropriate, variables were log-transformed. One-hot encoding was used for all factor variables prior to running RDA. The full list of all 54 climate and soil variables can be found in Table S2 and raw environmental data for each accession is provided in Table S12.

### 2.3 Spatial data

We represented spatial (non-genetic) structure in RDA using distance-based Moran Eigenvector Maps (dbMEMs; formally referred to as principle components of neighborhood matrices or PCNMs). Briefly, dbMEMs—much like principle components in a PCA—represent structure in a dataset with orthogonal axes. dbMEMs describe complex patterns of spatial structure in rectangular form suitable for use as a co-factor in downstream statistical analyses such as RDA (Borcard et al., 2018). Importantly, they are able to model both broad-and fine-scale patterns (Borcard et al., 2018; see Figure S1), both of which may affect the partitioning of genetic variation across large and highly heterogenous landscapes. dbMEMs were calculated using the quickMEM function included in Borcard et al. (2018). This function uses forward selection and permutation tests to identify the appropriate number of axes needed to explain patterns of spatial genetic variation, with the intention of preventing overfitting. We projected latitude and longitude data for each accession using the SIRGAS 2000 South America datum, which was then used as input to calculate dbMEMs.

### 2.4 Population genetic structure

Both model-based (*fastStructure*; Raj et al., 2014) and non-model based (PCA) methods were used to quantify genetic structure in the data. Markers in high linkage disequilibrium (r^2^ > 0.9) and those containing high levels of missing data (>5%) were pruned from the dataset prior to running PCA and *fastStructure* using the bcftools ‘prune’ plugin and vcftools, respectively (Li, 2011; (Danecek et al., 2011). The resulting dataset contained 7,317 high quality SNPs. PCA was implemented in the R (R Core Team, 2019) package adegenet (Jombart & Ahmed, 2011). Prior to running PCA, missing calls were imputed as the mean numeric value for each marker, to speed up computation. Model complexity in *fastStructure* was determined by minimizing the marginal likelihood of the data over values of K from one to ten. Clusters in the data were also identified using the K-means algorithm for all PCs by minimizing the Bayesian information criterion (BIC) across values of K from one to 100.

### 2.5 Partitioning total SNP variation

To evaluate the relative contribution of demographic history and the abiotic environment to explaining patterns of spatial genetic variation, the amount of genome-wide SNP diversity attributable to environment and geography was estimated using three methods: variance partitioning by redundancy analysis (RDA), generalized dissimilarity modeling (GDM), and structural equation modeling (SEM). All variance partitioning methods were performed with LD-filtered sites containing no missing data (6,830 SNPs, out of the 7,317 5% set above; Table S5). RDA is a form of constrained ordination first applied in community ecology that allows one to assess the explanatory power of multivariate predictors (in this case, environmental and geographic variables) for multivariate responses (genotypes; van den Wollenberg, 1977; Legendre & Legendre, 1998). RDA identifies linear combinations of explanatory variables (canonical axes) that best explain linear combinations of response variables. As with other forms of eigenanalysis, these canonical axes are orthogonal to each other. Here, these axes represent correlated SNPs that are also correlated with environment. Using multiple partially constrained models, it is possible to partition total genetic variation into that explained independently by climate, space, and their colinear portion (Borcard et al., 2018). All RDA analyses were performed using the R package vegan (Oksanen, 2018) and significance was assessed using 1,000 permutations of the genotype matrix. Since RDA requires no missing data, only markers containing no missing calls were included (6,830 SNPs). Distances between collection sites were represented using dbMEMs (described above). Because collinearity among environmental predictors can lead to inflation of variance components in RDA (Borcard et al., 2018), we performed forward variable selection using the forward.sel function in the R package adespatial to identify predictive and non-redundant environmental variables for partitioning.

In addition to RDA, structural equation modeling(SEM) was performed using the R package *lavaan* (Rosseel, 2012). SEM is a statistical framework similar to path analysis that has been applied to estimating the relative contributions of isolation-by-environment (IBE) and isolation-by-distance (IBD) to explaining spatial genetic variation (Wang et al., 2013). While powerful, this method is not able to estimate the independent contribution of the colinear portion of environment and space on explaining SNP variation, unlike RDA. For SEM, geographic distance was represented using Euclidean distance between accessions (calculated from projections using the SIRGAS 2000 datum). All 54 environmental variables were used to infer the latent environmental distance term (SEM accommodates correlated inputs; Wang et al., 2013). SEM was performed using the same set of markers as used for variance partitioning in RDA.

Lastly, generalized dissimilarity modeling (GDM; Ferrier et al., 2007) was applied to the SNP data. GDM is similar in principle to RDA and SEM but relies on different statistical methods. GDM uses nonlinear matrix regression to accommodate nonlinear statistical relationships that are often observed in ecological datasets; in contrast, RDA and SEM—as implemented here—do not model nonlinear relationships. GDM analyses were performed in R using the *gdm* package (Manion et al., 2018).

### 2.6 Importance of specific climate variables

Forward variable selection with RDA was used to infer which specific environmental variables may be most important for generating selective differences across the range of S. pimpinellifolium. We performed this procedure with the forward.sel function in adespatial, using 10,000 permutations to assess significance at each iteration and with RDA models both including and not including spatial structure as a covariate. The relative contribution of each individual variable to the constrained model was calculated as a sum of correlation coefficients across the first four RDA axes, weighted by the eigenvalue for each axis. The contribution of each variable to the full RDA model was then calculated as the ratio of this value and total genetic variance (Lasky et al., 2012).

### 2.7 Identifying outlier loci

RDA was also used to identify outlier loci showing especially strong relationships with multivariate environmental axes. The ability of RDA to identify loci responding to combinations of environmental predictors, instead of testing and treating each variable independently, is an advantage of RDA outlier analysis over classical mixed-model association methods. Moreover, because it simultaneously evaluates combinations of SNPs, RDA may also be better able to identify more subtle patterns environmental association involving multiple minimally diverged loci (Forester et al., 2018)—suggestive of polygenic adaptation—that can be obscured by traditional approaches. Here, outliers were defined as the SNPs having loadings along the first RDA axis +/− 4 SDs from the mean for that axis (Forester et al., 2018; Lasky et al., 2012). To reduce the potential for spurious associations due to shared ancestry, a partial RDA conditioned on dbMEMs was implemented. Instead of using the reduced set of markers with no missing calls for this analysis, we imputed SNPs at the markers with missing data in the full dataset (44, 064 SNPs) based on the mean allelic value at each site, and used this full dataset for outlier identification. Top candidate SNPs were investigated for their proximity to known gene models, using the ITAG 3.2 tomato genome annotation (Tomato Genome Consortium, 2012).

## 3 RESULTS

### 3.1 Genotype Coverage and Distribution

Our ddRAD procedure tagged 458,945 high coverage loci (effective per-sample coverage = 66.1x [sd = 36.7x]; average insert size = 139.5 bp; Table S3). The average number of loci seen in each individual was 60,044 (Table S4). Our genotyping pipeline identified 366,466 SNPs segregating within *S. pimpinellifolium* (Table S5). After stringent filtering for depth and missing data, this set of SNPs was reduced to 44,064. A second LD-filtered (r^2^ < 0.9) dataset was also generated for specific analyses where these data are preferable (see below); this dataset contained 17,358 SNPs (see Table S5 for details on genotype filtering). One variant occurred roughly every 18 kb. SNPs were enriched (75.6%) for sites in or nearby (within 5 kb) annotated gene models (9.4% within exons), suggesting that our enrichment based on *in silico* digestion was successful (determined by *SnpEff*; Cingolani et al., 2012). Of annotated SNPs in coding regions, 60.9% resulted in changes to amino acid composition (i.e., were non-synonymous).

### 3.2 Population structure follows a latitudinal gradient and defines regional genetic clusters

The first two axes of the multi-locus PCA based on 7,317 SNPs (LD pruned; <5% missing data) explained 18.03% and 6.53% of genetic variation, respectively. Minimizing BIC in K-means clustering identified five clusters in the data (Figure 2A & 2B), which followed a latitudinal gradient and roughly corresponded to variation along the first principal component axis. The model-based analysis conducted in *fastStructure* identified four groups in the data; these groups largely corresponded to the clusters identified using K-means, except that clusters four and five were not separated and instead were identified as a single group (Figure 2C; Figure S5). These patterns agree with other recent genomic analyses of population structure in this species (Lin et al., 2019). Varying degrees of admixture were inferred. The most heterogeneous individuals were found at the center and northern regions of the species range (clusters 1 [purple; Ecuador] and 3 [light blue; central Peru] in Figure 2), which matches general expectations for patterns of genetic diversity across species ranges (Vucetich & Waite, 2003). Similarly, patterns of heterozygosity (Figure S2) generally support a Northern Peruvian origin for *S. pimpinellifolium*, in agreement with previous marker-based analyses in this species (Rick et al., 1977; Zuriaga et al., 2009; Lin et al., 2019). Nonetheless, while some previous analyses have shown heterozygosity to specifically decrease in Northern Ecuador relative to Southern Ecuador and Northern Peru (corresponding to a unimodal cline in outcrossing centered on Northern Peru; Rick et al., 1977; Zuriaga et al., 2009), our data do not show a clear decline at the northern end of the species range, possibly because North Ecuador localities are more sparsely represented among our focal accessions and may not capture the northern tail of the cline in diversity.

**FIGURE 2.**
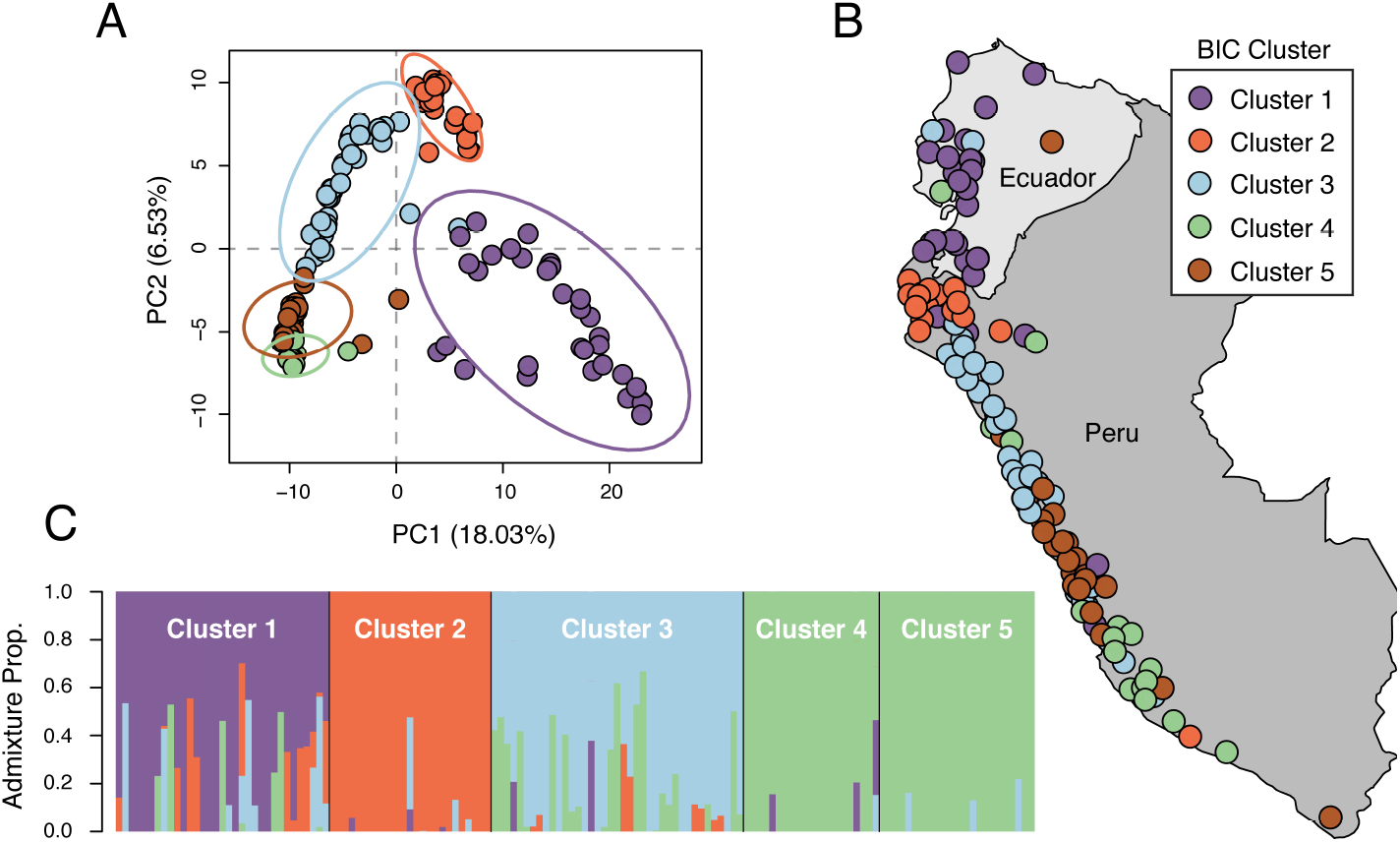
Genetic population structure in *S. pimpinellifolium* as revealed by principle components (PC) analysis (top panels) and *fastStructure* (bottom panel). (A) First two PC axes. Points are colored by membership in the five K-means clusters. (B) Accession collection sites colored by K-mean/BIC clin thuster membership. (C) Admixture proportions of all accessions identified with *fastStructure*, ordered by clusters identified in K-means to show correspondence between methods. BIC clusters 4 and 5 were identified as a single group using *fastStructure*.

### 3.3 RDA identifies climate and spatial variables that predict SNP variation

#### 3.3.1 Spatial variables

Forward selection of the dbMEM variables identified 10 axes as sufficient to explain geographic structure among accessions. Four of these axes described broad patterns of latitudinal structure (dbMEM1, 2, 3, and 4) and the remaining six described fine-scale patterns (dbMEM6, 7, 8, 9, 11 and 17; Figure S1). The axes describing fine-scale spatial structure cumulatively explained the most SNP variation among accessions (7.01%; Table 1). Broad-scale spatial variables explained a total of 5.42% of SNP variation (Table 1). dbMEM2 explained the most variation of any single variable (1.74%).

**TABLE 1.**
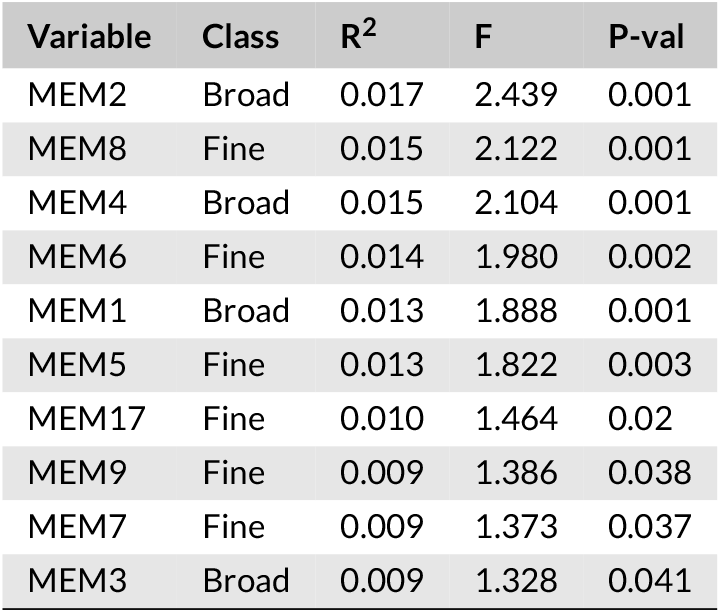
List of dbMEM variables and their contribution to SNP variation as determined by forward variable selection with RDA. Cumulatively more variation was explained by variables describing fine-scale geographic structure (0.070) than large-scale structure (0.054). Significance in forward selection was determined by permutation tests.

| Variable | Class | R<sup>2</sup> | F | P-val |
| --- | --- | --- | --- | --- |
| MEM2 | Broad | 0.017 | 2.439 | 0.001 |
| MEM8 | Fine | 0.015 | 2.122 | 0.001 |
| MEM4 | Broad | 0.015 | 2.104 | 0.001 |
| MEM6 | Fine | 0.014 | 1.980 | 0.002 |
| MEM1 | Broad | 0.013 | 1.888 | 0.001 |
| MEM5 | Fine | 0.013 | 1.822 | 0.003 |
| MEM17 | Fine | 0.010 | 1.464 | 0.02 |
| MEM9 | Fine | 0.009 | 1.386 | 0.038 |
| MEM7 | Fine | 0.009 | 1.373 | 0.037 |
| MEM3 | Broad | 0.009 | 1.328 | 0.041 |

#### 3.3.2 Climate variables

When analyzed without accounting for spatial structure using partially constrained ordination, we identified 14 abiotic variables as significantly predictive of genetic variation among accessions (Table 2). The most predictive variable was degree days above 18°C, however several variables contributed nearly equally to the RDA model. These include annual maximum solar radiation, average annual wind speed, isothermality (a measure of temperature variability), annual maximum wind speed, and variation in vapor pressure. In contrast, forward selection constrained on dbMEM spatial axes identified 6 environmental variables as predictive of genetic variation (Table 2); these variables represent gradients predictive of genetic variation after accounting for the proportion explained by geographic distance. Importantly, these will not include environmental gradients that are entirely colinear with space—including some that may contribute to clinal adaptation—and thus represent a conservative subset of the variables potentially contributing to geographical variation in selection. The contribution of these 6 gradients in RDA space is shown in Figure 3. The strongest predictor was the coefficient of monthly variation in vapor pressure, followed by precipitation seasonality, soil texture, annual maximum solar radiation, maximum potential evapotranspiration, and minimum potential evapotranspiration (Table 2).

**TABLE 2.** Top environmental variables and their contribution to spatial SNP variation as determined by forward variable selection with RDA. Results from both the full RDA model selection (not conditioned on space) and partial RDA model selection (conditioned on space) are reported.

| RDA (not constrained on space) |  | RDA (constrained on space) |  |
| --- | --- | --- | --- |
| Variable | Contribution to RDA model | Variable | Contribution to RDA model |
| Degree days above 18C | 13.60 | CV vapor pressure | 2.76 |
| Annual max solar radiation | 13.45 | Prec. seasonality | 2.43 |
| Annual average wind speed | 13.45 | Soil texture | 2.25 |
| Isothermality | 13.44 | Annual max solar radiation | 2.23 |
| Annual max wind speed | 13.33 | Max potential evapotransp. | 2.16 |
| CV vapor pressure | 13.31 | Min potential evapotransp. | 1.64 |
| Temperature seasonality | 13.17 |  |  |
| Extreme min temperature (30 yr) | 13.10 |  |  |
| Max soil water stress | 11.55 |  |  |
| Climatic moisture deficit | 10.63 |  |  |
| Annual temperature range | 9.44 |  |  |
| Mean diurnal range | 8.02 |  |  |
| Max potential evapotransp. | 6.05 |  |  |
| CV wind speed | 3.21 |  |  |

**FIGURE 3.**
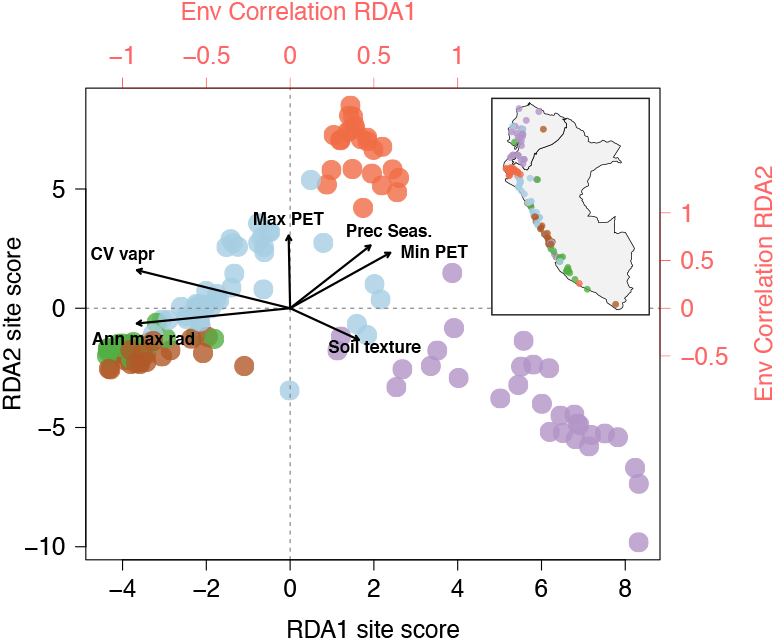
Biplot of site (accession) scores for the first two RDA axes, constraining on space with dbMEMs. Points are colored to correspond with genetic clusters identified in PCA (shown in inset, from Figure 2). Top and right axes (red) show the correlation of each environmental predictor with RDA axes 1 and 2, respectively.

The correlations of these six variables with the first two RDA axes (loadings in Figure 3) suggests that the contribution of each variable to SNP variation differs geographically across the species range, so that the environmental predictors most influential in driving SNP divergence between genetic/geographic clusters varies depending upon the specific clusters being compared. To test for the presence of regional climatic differences that may contribute to divergent selection, we studied how each environmental predictor varied by genetic cluster (Figure S3). Significant differences among genetic clusters were identified for all 6 predictors based on ANOVA tests (P < 1 × 10^−8^ for all tests), suggesting that each regional cluster occupies a more or less unique abiotic habitat. This finding is recapitulated when breaking down all 58 environmental variables into orthogonal components (Figure S4); the first two components of climatic variation separate accessions by region.

### 3.4 Regional climatic differences explain more SNP variation than geography

#### 3.4.1 Variance partitioning by RDA

Using RDA, the abiotic environment (14 variables identified through forward selection) and spatial structure (10 dbMEMs identified through forward selection) explained 22% of SNP variation among accessions. The majority of this (17%) was attributable to the colinear portion of environment + space (Figure 4). This fraction could represent either the contribution of clinal environmental factors or latent spatial population structure that is not captured by the dbMEMs. Considering the independent effects of environment and space, the environment alone explained more SNP variation (3%) than space alone (2%; Figure 4). The observed proportions of variation explained by climate and space were greater than the proportions explained in all of 1000 permuted data sets (P < 0.001).

**FIGURE 4.**
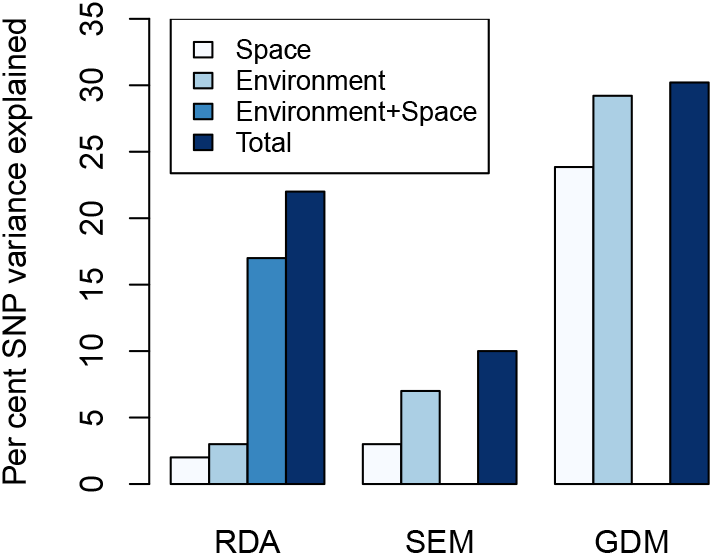
Summary of the three variance partitioning methods. Note that SEM and GDM are unable to quantify the independent contribution of environment + space.

#### 3.4.2 Structural equation modeling (SEM)

Regression components of environmental distance and geographic distance on genetic distance identified using SEM were 0.788 (SE = 0.044) and 0.174 (SE = 0.012), respectively (Table S6). This corresponds to 7% (6.4% 7.7%) and 3% (2.6% 3.5%) of total genetic variance being attributable to environmental distance and geographic distance between accessions, respectively, which is consistent with our findings from RDA (Figure 4). Only the environmental regression component was higher than those obtained from all of 1000 permuted data sets (P < 0.001). As with RDA, this suggests that environmental differences between accessions are shaping the distribution of genetic variation to a larger degree than spatial isolation.

#### 3.4.3 Generalized dissimilarity modeling (GDM)

The percent deviance in genetic variation among accessions explained by the full (environment + space) GDM model was 30.2% (Figure 4; see Table S7 for full model fit and Table S8 & S9 for variable contributions). 29.2% and 23.9% of deviance was explained by models considering only environmental and geographic effects on genetic variation between accessions, respectively (Figure 4). All of these models were significant (P < 0.001) based on permutation tests. These results agree with RDA and SEM; a larger fraction of SNP variation is attributable to environmental variation than to spatial isolation.

### 3.5 RDA identifies outlier loci associated with multivariate climate

We observed significant variation among SNPs in the degree to which they were associated with the abiotic environment, as measured by their loadings along the first RDA axis (Figure 5). We identified candidate loci for local adaptation by examining SNPs displaying loadings along the first RDA axis +/− 4 standard deviations from the mean (Forester et al., 2018; Lasky et al., 2012). This identified 252 SNPs showing particularly strong associations with multivariate climate. In the model containing spatial structure as a covariate, the strongest signal of association spanned 23.5 Mbp on chromosome four (Figure 5). This region contained 333 annotated genes. The top associated SNP in this region was 30 bp upstream of a 60S ribosomal subunit gene (Table 3). Other significant associations in this region include intergenic substitutions proximal to DNA glycosylase, exocyst complex component 7, and cellulose synthase; a non-synonymous substitution in MYB transcription factor 73; and intronic substitutions in sterol glusoyltransferase 4 (SGT4), nuclease S1, BY-2 kinesin protein 5, RNA helicase DEAH-Box 13, and translation initiation factor 3, among others (Table 3; Table S11). No outliers identified in the analysis without spatial covariates were also present in the space + environment constrained RDA analysis. This is consistent with our general observation of a strong degree of collinearity between genetic and geographical variation in this species (Figure 4; Table S10), and reiterates the need to include spatial structure when evaluating the independent contribution of environmental factors to SNP variation.

**TABLE 3.** Annotated gene models located within 10kb of the top 15 outlier SNPs along the first RDA axis, after accounting for spatial structure. We also do not report SNPs within 10kb of SNPs with higher loadings. The full list of outlier SNPs and their annotations can be found in table S8.

| Chr. | SNP Position | Locus | SNP Category | Distance from Locus (bp) | RDA1 Loading | Locus Description |
| --- | --- | --- | --- | --- | --- | --- |
| 4 | 46907724 | Solyc04g050390 | Intergenic | 30 | 0.015 | 60S ribosomal subunit |
| 4 | 37812488 | Solyc04g047830 | Intergenic | 4389 | 0.013 | DNA glycosylase |
| 4 | 44823446 | Solyc04g049930 | Missense | 0 | 0.013 | Unknown protein |
| 4 | 49678371 | Solyc04g051150 | Intron | 0 | 0.013 | Sterol glucosyl transferase 4 (SGT4) |
| 3 | 66381751 | Solyc03g115060 | Intergenic | 66 | 0.012 | Exocyst complex component 7 |
| 5 | 3609439 | Solyc05g009440 | Intron | 0 | 0.012 | Nuclease S1 |
| 1 | 88554548 | Solyc01g098080 | Intron | 0 | 0.011 | BY-2 kinesin-like protein 5 |
| 4 | 45372222 | Solyc04g050080 | Missense | 0 | 0.011 | MYB transcription factor 73 |
| 8 | 26166364 | Solyc08g041710 | Intron | 0 | 0.011 | Transmembrane protein |
| 6 | 39445574 | Solyc06g062360 | Intron | 0 | 0.011 | Syntaxin-like protein |
| 11 | 417966 | Solyc11g005560 | Intergenic | 658 | 0.011 | Cellulose synthase |
| 8 | 27570643 | Solyc08g023500 | Intron | 0 | 0.011 | Metallohydrolase/oxioreductase |
| 4 | 5709089 | Solyc04g015490 | 3' UTR | 0 | 0.011 | Magnesium chelatase subunit D |
| 4 | 45599110 | Solyc04g050150 | Intron | 0 | 0.011 | RNA helicase DEAH-Box 13 |
| 8 | 23509033 | Solyc08g042140 | Intron | 0 | 0.011 | Translation initiation factor 3 subunit |

**FIGURE 5.**
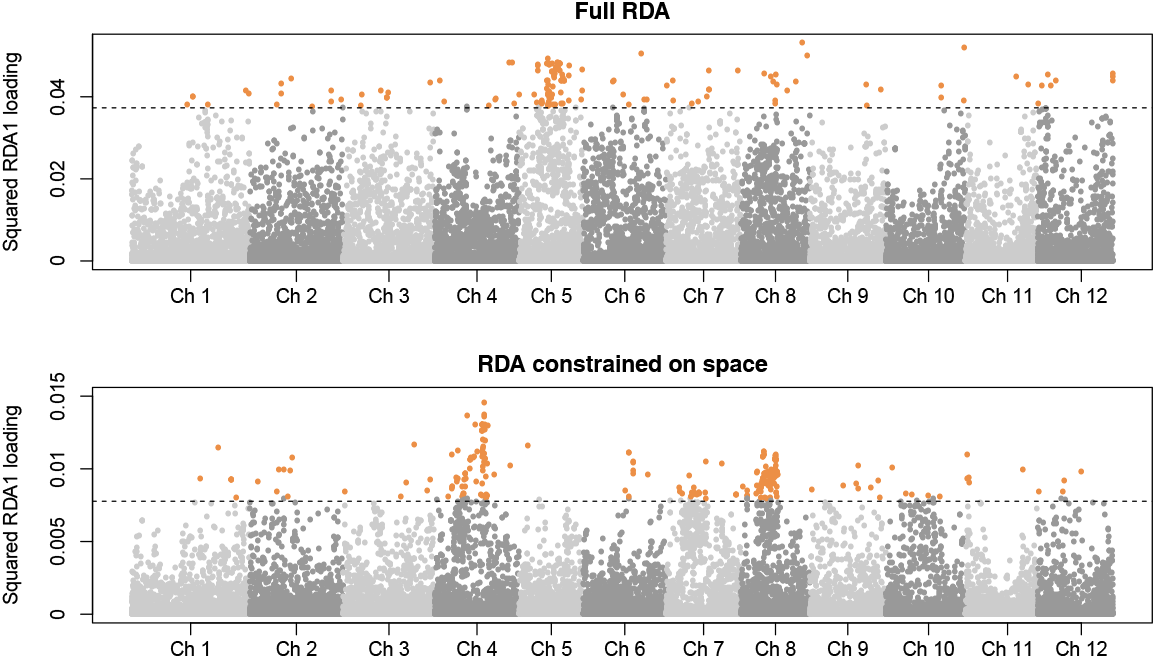
Values of squared RDA1 SNP loadings throughout the genome. Top panel: SNP loadings on the first RDA axis of raw SNP variation. Bottom panel: SNP loadings on the first RDA axis accounting for spatial structure among accessions. Note the change in y-axis scale between the two panels. Dashed lines represent +/− 4 SD cutoff used to identify outliers.

## 4 DISCUSSION

Local adaptation to the abiotic environment is often thought to play a major role in diversification both within and between species, however its contribution relative to other evolutionary forces is rarely quantified. In this study, we investigated the relationship of genome-wide SNPs with fine-scale climatic and spatial data among 140 accessions of the ecologically diverse crop wild relative *S. pimpinellifolium*. Our study affirms previous work demonstrating broad correlations between genetic and climatic structure in this system (Blanca et al., 2012, 2015; Lin et al., 2019; Zuriaga et al., 2012), and builds beyond this to quantify the contributions of individual environmental variables to patterns of genomic variation and dissect the genetic targets of these putative selective gradients in a multivariate statistical context. We found that the abiotic environment—as described by climate and soil characteristics—explained more SNP variation than spatial structure (Figure 4), demonstrating that our evaluated environmental factors are more influential in shaping the geographic distributions of SNPs in this species, compared to spatial isolation. Both GDM and SEM—two methods which have also been applied to quantifying the contributions of space and environment to patterns of genetic variation—agree with our RDA results. From this we conclude that ecological differences among populations are closely associated with the maintenance of contemporary genetic variation across space and that local adaptation is likely a major driver of these patterns.

Our RDA analysis identified that a large (17%) portion of SNP variation among accessions was explainable by the colinear fraction of environment and space (Figure 4). This fraction could represent the contribution of clinal environmental forces that are highly correlated with space (e.g., latitudinal gradients in temperature). Clinal gradients are considered to be important drivers of adaptive evolution in many species (Adrion et al., 2015; Kooyers et al., 2015), and across the range of *S. pimpinellifolium*, many environmental variables show strong geographic trends and are correlated with both genetic (i.e., PCs) and spatial (i.e., dbMEMs) structure (Table S10). Previous studies in *S. pimpinellifolium* have demonstrated strong correlations between environmental and genetic distances among accessions of this species, largely associated with latitudinal gradients (Blanca et al., 2015; Zuriaga et al., 2009). Accordingly, when we did not correct for spatial structure with constrained ordination, a highly clinal variable—degree days above 18C—explained the most total SNP variation (Table 2). Although geographic clines have historically been considered important for understanding the genetics of adaptation (Adrion et al., 2015; Endler, 1977, 1986), disentangling geographically structured variation from adaptive variation remains a major challenge—as evidenced here—that will likely require novel statistical methods and experimental studies to resolve. In *S. pimpinellifolium*, further population sampling across clinal gradients coupled with *in situ* common garden experiments could be one way of disentangling the independent effects of space and climate in these cases.

Despite the substantial collinearity we observed between space and environment, our analysis discerned several axes of spatial variation that uniquely explained SNP variation. The second spatial dbMEM variable explained the largest portion of SNP variation among accessions (Table 1). This variable describes broad patterns of spatial structure and separates accessions from the center of the range (northern Peru) from those at the northern and southern extremes (Figure S1). This finding is consistent with our genetic analysis of population structure—as well as previous spatial genetic data (Rick et al., 1977; Zuriaga et al., 2009; Lin et al., 2019)—which demonstrate a strong latitudinal pattern of genetic relatedness. Interestingly, while the second dbMEM axis explained the most variance of any single spatial variable, cumulatively more variance was explained by fine-scale dbMEM axes (dbMEM5, 6, 7, 8, 9, & 17; cumulative R^2^ = 0.0701). Such a pattern may be driven by the geographic landscape of the Andes; large and patchy altitudinal clines (sea level to > 6500 m) may limit local dispersal between adjacent populations, amplifying genetic differentiation over small spatial scales.

Apart from spatial variation associated with SNPs, our analysis also identified specific environmental factors that uniquely explained additional SNP variation. Temporal fluctuations in precipitation (seasonality) and vapor pressure (the coefficient of monthly variation) explained the most genome-wide SNP variation when we simultaneously considered the effects of spatial structure in our RDA analysis (Table 2). This reinforces some previous empirical findings—especially the role of precipitation seasonality as a key factor—but also points to additional environmental forces as potential abiotic selective agents in this species. Precipitation was previously implicated in driving ecological adaptation in both this and other tomato species (Nakazato et al., 2008, 2010). Specifically, among accessions of *S. pimpinellifolium*, variation in days to wilting (a measure of drought tolerance) is strongly associated with differences in annual precipitation between coastal and highland regions (Nakazato et al., 2008), consistent with selection being imposed by this environmental axis. Interestingly, our findings at the intraspecific scale are also similar to previous inferences of important ecological drivers among wild tomato species. Distribution models have identified precipitation and temperature as the major determinants of differences in contemporary species ranges across the wild tomato clade (Nakazato et al., 2010). Moreover, these analyses indicate that precipitation seasonality in particular is important for determining the allopatric distributions of *S. pimpinellifolium* and its sister taxon *S. lycopersicum* var. *cerasiforme*.

In contrast to precipitation, the importance of vapor pressure as a selective agent within or among wild tomato species has not been evaluated, although vapor pressure’s role in modulating plant-water relations has been well studied in domesticated tomato (Guichard et al., 2005; Zhang et al., 2017). Deficits in vapor pressure between the mesophyll and external leaf surface promotes transpiration (Zhang et al., 2017), and if not met with additional water uptake from the soil, large differentials in vapor pressure will lead to reduced growth and desiccation. Our findings from RDA suggest that regional differences in vapor pressure are a strong selective force in *S. pimpinellifolium* (Table 2). The importance of variation in vapor pressure—and not a measure of annual or monthly mean values—may point to a role for specific physiological mechanisms which maximize fitness during acute fluctuations in evaporative demand (e.g., rapid stomatal closure responses, induction of root growth, or other strategies for increasing water use efficiency under temporary stress). This hypothesis is testable in the future with direct manipulative experiments across accessions that are predicted to differ at specific SNPs associated with this axis of the environment.

While variation in vapor pressure and precipitation were the most predictive of genomic variation across the range as a whole (Table 2), regional environmental differences and our RDA analysis suggest that different specific factors are most influential in different regions of the species range (Figure 3; Figure S3). Genetic clusters we identify within *S. pimpinellifolium* are, in general, significantly diverged from each other for their environments (Figure S3; Figure S4). For example, coastal accessions (genetic clusters 1, 4, and partially 3) are exposed to higher levels of solar radiation compared to the rest of the range (Figure S3, panel 3); northern Peruvian accessions (cluster 2) experience highly seasonal precipitation (Figure S3; panel 2); and Ecuadorian accessions (cluster 5) experience high soil type diversity (i.e., all four soil types that found throughout the species range are also present in this region; Figure S3, panel 6). Together with our RDA results—which identified these and other variables as significantly predictive of SNP variation independent of spatial population structure—these patterns of environmental variation are consistent with regional differences in the primary agents of ecological selection. Unlike studies of genetic variation across precipitation gradients (see above), equivalent analyses of genetic variation for tolerance to these environmental factors (i.e., solar radiation, soil texture, and potential evapotranspiration) is currently limited. Further common garden manipulation experiments would be helpful to understand the exact role these forces have in shaping local climatic adaptation in *S. pimpinellifolium*. Nonetheless, these observations broadly mirror comparative findings from among wild tomatoes—that detect clear niche differentiation between species (Nakazato et al., 2010)—by uncovering similar niche variation intraspecifically at the regional level.

In addition to these findings, and just as with other genetic (Lasky et al., 2012, 2016) and community ecological studies using similar methods (Borcard et al., 2018), a large portion of variance remained unexplained. This is likely due to the influence of several factors that cannot be fully addressed in this type of experiment. First, although we included a large set of environmental variables in our analyses, many other unmeasured ecological forces could also be acting; most importantly, this includes biotic interactions that are likely only partially reflected in the abiotic factors considered here. Second, axes of population structure may not be fully explained by spatial dbMEM variables, potentially due to unmodeled geographic heterogeneity present in the Andes. Third, other evolutionary forces that maintain local allelic diversity (e.g., balancing selection) may weaken signals of SNP-environment associations. Lastly, as implemented here, both RDA and SEM only model linear associations between climate/space and SNPs (Borcard et al., 2018), and thus will not capture non-linear statistical relationships if they exist. Interestingly, our GDM analysis—which does model nonlinear associations—explained more total SNP variation than the other two methods (Figure 4), suggesting the presence of nonlinear SNP-environment relationships across the range of *S. pimpinellifolium*. Explicitly modeling and estimating the contribution of nonlinear terms to patterns of SNP-environment associations may be crucial to understanding local adaptation in the context of multivariate environments, but such methods are currently limited (Rellstab et al., 2015).

Our SNP outlier analysis allowed us to map the location of genomic regions showing strong associations with major axes of the environment and identify candidate loci potentially involved in local adaptation (Figure 5). In contrast to traditional scans that treat each environmental variable independently (e.g., Fournier-Level et al., 2011; Hancock et al., 2011), our analysis considered how loci are associated with linear combinations of environmental gradients. This framework may be more appropriate for identifying ecologically relevant loci; rarely can a single environmental factor be considered the only contributor to selection. We identified 252 signals of putatively adaptive loci in *S. pimpinellifolium* using genomic scans based on RDA loadings. Of foremost interest are two regions containing known regulators of abiotic stress tolerance: Solyc04g051150—sterol glucosyl transferase 4 (SGT4)—and Solyc04g050080—MYB transcription factor 73 (Table 3). SGT genes have been shown to have functional roles in abiotic stress tolerance in several species (Griebel & Zeier, 2010; Hugly et al., 1990; Kumar et al., 2015; Senthil-Kumar et al., 2013). Expression of SGT4 and its paralog SGT2 is rapidly induced upon exposure to cold and salt stress in domesticated tomato (Ramirez-Estrada et al., 2017). In Arabidopsis thaliana SGT knock-out mutants, plants show reduced tolerance to several abiotic stressors, including cold and salinity (Mishra et al., 2015). MYB 73 acts as a salt stress-specific transcription factor and modulates whole-plant responses to salinity in A. thaliana (Kim et al., 2013). Expression profiles of domesticated tomato under hormone-induced stress suggest a general role for MYB transcription factors in responding to abiotic cues (Li et al., 2016), although the specific function of MYB 73 has not been evaluated in Solanum. While genotypic variance in salt-stress tolerance has been documented in *S. pimpinellifolium* (Rao et al., 2013)—suggesting that functional variation in salt-response loci may be prevalent—our analyses were not able to evaluate the contribution or association off this factor to genome-wide SNP variation, since fine-scale data on soil salinity in South America is not yet available. Nonetheless, these scans provide new information on the location of loci relevant for adaptive responses to abiotic stress, generating prime candidates for future functional investigation, as well as for breeding into domesticated lines.

Overall, our study adds to an emerging landscape genomics literature applying an ensemble of multivariate methods to dissect the demographic and environmental features shaping patterns of genomic differentiation. Quantifying the influence of local adaptation to climatic variation has profound importance for predicting how species will respond to future climates, for the breeding of resilient cultivars, and for broadly understanding selection as a mechanism of diversification. Here we generate a strong case that environmentally imposed selection plays a key role in genetic differentiation within a widespread and diverse crop wild relative, and distinguish the climatic factors most likely responsible. All three multivariate methods (RDA, SEM, and GDM) identified that climatic variation among our 140 *S. pimpinellifolium* accessions explained more genomic variation than does geographic distance, with temporal variation in precipitation and evaporative demand explaining the most SNP variation overall. Our analysis also identified significant geographical differences in other key climatic factors that suggest additional intraspecific niche differentiation between geographic regions. These results complement comparative findings from the wild tomato clade (Nakazato et al., 2010) indicating that spatially varying abiotic selection may be an important source of genetic variability in this species, but also demonstrate the additional qualitative and quantitative insight that can be gained by applying a lens of landscape genomics. This additional power includes uncovering genomic regions with strong environmental associations—potential genomic targets of ecological selection—that can be passed to further studies to understand their exact genetic and phenotypic contributions, for example, by incorporating SNP-environment associations into breeding value prediction frameworks (Lasky et al., 2015; Yoder et al., 2012). Together, our findings add to a growing literature demonstrating a pervasive role of selection in shaping patterns of genomic differentiation, and the likely multilocus architecture that underlies adaptive differentiation, using landscape genomics. Additional studies in a broader range of species and ecological contexts will be critical for assessing additional generalities about the demographic and environmental features most influential in defining patterns of genetic variation.

## Ackowledgements

The authors thank the UC Davis Tomato Genetic Resource Center for providing access to the *S. pimpinellifolium* germplasm used in this study, the Indiana University Center for Genomics and Bioinformatics for library preparation and sequencing, the Indiana University greenhouse staff, and three anonymous reviewers for comments that improved a previous version of this manuscript. This research was supported by National Science Foundation award DEB1136707 to LCM.

## Author Contributions

MJSG and LCM designed the experiments; MJSG performed the greenhouse and molecular work and conducted the bioinformatic analyses; MJSG and LCM wrote the manuscript.

## REFERENCES

Adrion, J. R., Hahn, M. W. and Cooper, B. S. (2015) Revisiting classic clines in Drosophila melanogaster in the age of genomics. Trends in Genetics, 31, 434–444. URL: http://dx.doi.org/10.1016/j.tig.2015.05.006.

Andrews, S. (2010) FastQC: a quality control tool for high throughput sequence data.

Blanca, J., Canizares, J., Cordero, L., Pascual, L., Diez, M. and Nuez, F. (2012) Variation Revealed by SNP Genotyping and Morphology Provides Insight into the Origin of the Tomato. PLoS One, 7, e48198.

Blanca, J., Montero-Pau, J., Sauvage, C., Bauchet, G., Illa, E., Diez, N., Francis, D., Causse, M., van der Knaap, E. and Canizares, J. (2015) Genomic variation in tomato, from wild ancestors to contemporary breeding accessions. BMC Genomics, 16, 257.

Bolarin, M., Fernandez, F., Cruz, V. and Cuartero, J. (1991) Salinity tolerance in four wild tomato species using vegetative yieldsalinity response curves. Journal of the American Society for Horticultural Science, 116, 286–290.

Borcard, D., Gillet, F. and Legendre, P. (2018) Numerical Ecology with R. Springer International, 2 edn.

Catchen, J., Hohenlohe, P., Bassham, S., Amores, A. and Cresko, W. (2013) Stacks: an analysis tool set for population genomics. Molecular Ecology, 22, 3124–3140.

Chen, S., Zhou, Y. and Gu, J. (2018) fastp: an ultra-fast all-in-one FASTQ preprocessor. Bioinformatics, 34, i884–i890.

Chitwood, D., Headland, L., Filiault, D., Kumar, R., Jimenez-Gomeez, J., Schrager, A., Park, D., Peng, J., Sinha, N. and Maloof, J. (2012a) Native environment modulates leaf size and response to simulated foliar shade across wild tomato species. PLoS One, 7, e29570.

Chitwood, D., Headland, L., Kumar, R., Peng, J., Maloof, J. and Sinha, N. (2012b) The developmental trajectory of leaflet morphology in wild tomato species. Plant Physiology, 158, 1230–1240.

Cingolani, P., Platts, A., Wang, L., Coon, M., Nguyem, T., Wang, L., Landm S.J, Lu, X. and Ruden, D. (2012) A program for annotating and predicting the effects of single nucleotide polymorphisms, SnpEff: SNPs in the genome of Drosophila melanogaster strain w1118; iso-2; iso-3. Fly, 6, 1–13.

Consortium, T. G. (2012) The tomato genome sequence provides insights into fleshy fruit evolution. Nature, 485, 635–641.

Cuartero, J., Yeo, A. and Flowers, T. (1992) Selection of donors for salt-tolerance in tomato using physiological traits. New Phytologist, 121, 63–69.

Danecek, P., Auton, A., Abecasis, G., Albers, C. A., Banks, E., DePristo, M. A., Handsaker, R. E., Lunter, G., Marth, G. T., Sherry, S. T., McVean, G., Durbin, R. and Group,. G. P. A. (2011) The variant call format and VCFtools. Bioinformatics, 27, 2156–2158. URL: https://doi.org/10.1093/bioinformatics/btr330.

Des Marais, D. and Juenger, T. (2010) Pleiotropy, plasticity, and the evolution of plant abiotic stress tolerance. Annals of the New York Academy of Sciences, 1206, 56–79.

Des Marais, D., McKay, J., Richards, J., Sen, S., Wayne, T. and Juenger, T. (2012) Physiological genomics of response to soil drying in diverse Arabidopsis acceessions. The Plant cell, 24, 893–914.

Endler, J. (1977) Geographic Variation, Speciation, and Clines. Princeton, NJ.: Princeton University Press.

Endler, J. (1986) Natural Selection in the Wild. Princeton, NJ.: Princeton University Press.

Ferrier, S., Manion, G., Elith, J. and Richardson, K. (2007) Using generalized dissimilarity modelling to analyse and predict patterns of beta diversity in regional biodiversity assessment. Diversity and Distributions, 13, 252–264.

Fick, S. and Hijmans, R. (2017) Worldclim 2: New 1-km spatial resolution climate surfaces for global land areas. Intenational Jounrnal of Climatology.

Foolad, M. and Yin, G. (1997) Genetic potential for salt tolerance during germination in Lycopersicon species. HortScience, 32, 296–300.

Foolad, M. R. (2007) Current status of breeding tomatoes for salt and drought tolerance. In. Advances in Molecular Breeding Toward Drought and Salt Tolerant Crops (eds. M. Jenks, P. Hasegawa and S. Jain). Springer.

Forester, B., Lasky, J., Wagner, H. and Urbal, D. (2018) Comparing methods for detecting multilocus adaptation with multivariate genotype-environment associations. Molecular Ecology, 27, 2215–2233.

Fournier-Level, A., Korte, A., Cooper, M., Nordborg, M., Schmitt, J. and Wilczek, A. (2011) A map of local adaptation in Arabidopsis thaliana. Science, 334, 86–89.

Griebel, T. and Zeier, J. (2010) A role for beta-sitosterol to stigmasterol conversion in plant-pathogen interactions. Plant Journal, 63, 254–268.

Guichard, S., Gary, C., Leonardi, C. and Bertin, N. (2005) Analysis of Growth and Water Relations of Tomato Fruits in Relation to Air Vapor Pressure Deficit and Plant Fruit Load. Journal Of Plant Growth Regulation, 24, 201.

Hamann, A., Wang, T., Spittlehouse, D. and Murdock, T. (2013) A comprehensive, high-resolution database of historical and projected climate surfaces for western North America. Bulletin of the American Meteorological Society, 94, 1307–1309.

Hancock, A., Brachi, B., Faure, N., Horton, M., Jarymowycz, L., Sperone, G., Toomajian, C., Roux, F. and Bergelson, J. (2011) Adaptation to Climate Across the Arabidopsis thaliana Genome. Science, 334, 83–86.

Hengl, T., Mendes de Jesus, J., Heuvelink, G., Ruiperez Gonzalez, M., Kilibarda, M., Blagotic, A., Shangguan, W., Wrightt, M., Geng, X., Bauer-Marschallinger, B., Guevara, M., Vargas, R., MacMillan, R., Batjes, N., Leenaars, J., Ribeiro, E., Wheeler, I., Mantel, S. and Kempen, B. (2017) Soil-Grids250m: Global gridded soil information based on machine learning. PLoS One, 12, e0169748.

Hodgins-Davis, A. and Towsend, J. (2009) Evolving gene expression: from G to E to G x E. Trends in Ecology & Evolution, 24, 649–658.

Hugly, S., McCourt, P., Browse, J., Patterson, G. and Somrville, C. (1990) A Chilling Sensitive Mutant of Arabidopsis with Altered Steryl-Ester Metabolism. Plant Physiology, 93, 1053–1062.

Jiang, J., Ma, S., Ye, N., Jiang, M., Cao, J. and Zhang, J. (2017) WRKY transcription factors in plant responses to stresses. Journal of Integrative Plant Biology, 59, 86–101.

Jissoudis, C., Sunarti, S., van de Wiel, C., Visser, R., van der Linden, C. and Bai, Y. (2016) Responses to combined abiotic and biotic stress in tomato are governed by stress intensity and resistance mechanism. Journal of Experimental Botany, 67, 5119–5132.

Jombart, T. and Ahmed, I. (2011) Adegenet 1.3-1: new tools for the analysis of genome-wide SNP data. Bioinformatics2, 27, 3070–3071.

Kim, J., Nguyen, N., Jeong, C., Nguyen, N., Hong, S. and Lee, H. (2013) Loss of the R2R3 MYB, AtMyb73, causes hyperinduction of the SOS1 and SOS3 genes in response to high salinity in Arabidopsis. Journal of Plant Physiology, 170, 1461–1465.

Klepikova, A., Kasianov, A., Gerasimov, E., Logacheva, M. and Penin, A. (2016) A high resolution map of the Arabidopsis thaliana developmental transcriptome based on RNA-seq profiling. Plant Journal, 88, 1058–1070.

Kooyers, N., Greenlee, A., Colicchio, J., Oh, M. and Blackman, B. (2015) Replicate altitudinal clines reveal that evolutionary flexibility underlies adaptation to drought stress in annual Mimulus guttatus. New Phytologist, 206, 152–165.

Kumar, M., Ali, K., Dahuja, A. and Tyagi, A. (2015) Role of phytosterols in drought stress tolerance in rice. Plant Physiology and Biochemistry, 96, 83–89.

Lasky, J., Des Marais, D., McKay, J., Richards, J., Juenger, T. and Keitt, T. (2012) Characterizing genomic variation of Arabidopsis thaliana: the roles of geography and climate. Molecular Ecology, 21, 5512–5529.

Lasky, J., Upadhyaya, H., Ramu, P., Deshhhpande, S., Hash, C., Bonnette, J., Juenger, T., Hyma, K., Acharya, C., Mitchell, S., Buckler, E., Brenton, Z., Kresovich, S. and Morris, G. (2015) Genome-environment associations in sorghum landraces predict adaptive traits. Science Advances, 1, e1400218.

Lee, C. and Mitchell-Olds, T. (2011) Quantifying effects of environmental and geographical factors on patterns of genetic differentiation. Molecular Ecology, 20, 4631–4642.

Legendre, P. and Legendre, L. (1998) Numerical Ecology. New York: Elsevier, 2 edn.

Li, H. (2011) A statistical framework for SNP calling, mutation discovery, association mapping and population genetical parameter estimation from sequencing data. Bioinformatics, 27, 2987–2993.

Li, H. and Durbin, R. (2009) Fast and accurate short read alignment with Burrows-Wheeler Transform. Bioinformatics, 25, 1754–1760.

Li, Z., Peng, R., Tian, Y., Han, H., Xu, J. and Yao, Q. (2016) Genome-Wide Identification and Analysis of the MYB Transcription Factor Superfamily in Solanum lycopersicum. Plant Cell and Physiology, 57, 1657–1677.

Lin, Y., Liu, C. and Chen, K. (2019) Assessment of Genetic Differentiation and Linkage Disequilibrium in Solanum pimpinellifolium Using Genome-Wide High-Density SNP Markers. G3, 9, 1497–1505.

Manel, S., Poncet, B., Legendre, P., Gugerli, F. and Holderegger, R. (2010) Common factors drive adaptive genetic variation at different spatial scales in Arabis alpina. Molecular Ecology, 19, 3824–3835.

Manion, G., Lisk, M., Ferrier, S., Nieto-Lugilde, D., Mokany, K. and Fitzpatrick, M. (2018) gdm: Generalized Dissimilarity Modeling.

Mishra, M., Singh, G., Tiwari, S., Singh, R., Kumari, N. and Misrra, P. (2015) Characterization of Arabidopsis sterol glycosyltransferase TTG15/UGT80B1 role during freeze and heat stress. Plant Signaling & Behavior, 10, e1075682.

Nakazato, T., Bogonovich, M. and Moyle, L. (2008) Environmental factors predict adaptive phenotypic differentiation within and between two wild andean tomatoes. Evolution, 62, 774–792.

Nakazato, T., Warren, D. and Moyle, L. (2010) Ecological and geographic modes of species divergence in wild tomatoes. American Journal of Botany, 97, 680–693.

Oksanen, J., Blanchet, F., Kindt, R., Legendre, P., Minchin, P., O’Hara, R., Simpson, G., Solymos, P., Stevens, M., Wagnerr, H. and Oksanen, M. (2013) Package ‘vegan’. Community ecology package, version 2.9.

Pedley, K. and Martin, G. (2003) Molecular basis of Pto-mediated resistance to bacterial speck disease in tomato. Annual Review of Phytopathology, 41, 215–243.

Pitblado, R. and Kerr, E. (1979) A source of resistance to bactereial speck–Pseudomonas tomato. Tomato Genetics Cooperative, 29, 30.

Raj, A., Stephen, M. and Pritchard, J. (2014) fastSTRUCTURE: Variational Inference of Population Structure in Large SNP Data Sets. Genetics2, 197, 573–589.

Ramirez-Estrada, K., Castillo, N., Lara, J., Arro, M., Boronat, A., Ferrer, A. and Altabella, T. (2017) Tomato UDP-Glucose Sterol Glycosyltransferases: A Family of Developmental and Stress Regulated Genes that Encode Cytosolic and Membrane-Associated Forms of the Enzyme. Frontiers in Plant Science, 8.

Rao, E., Kadirvel, P., Symonds, R. and Ebert, A. (2013) Relationship between survival and yield related traits in Solanum pimpinellifolium under salt stress. Euphytica, 190, 215–228.

Razali, R., Bougouffa, S., Morton, M., Lightfoot, D., Alam, I., Essack, M., Arold, S., Kamau, A., Schmockel, S., Pailles, Y., Shahid, M., Michell, C., Salim, A., Ho, Y., Tester, M., Bajic, V. and Negrao, S. (2018) The Genome Sequence of the Wild Tomato Solanum pimpinellifolium Provides Insights Into Salinity Tolerance. Frontiers in Plant Science, 9.

Rellstab, C., Gugerli, F., Eckert, A., Hancock, A. and Holderegger, R. (2015) A practical guide to environmental association analysis in landscape genomics. Molecular Ecology, 24, 4348–4370.

Rick, C., Fobes, J. and Holle, M. (1977) Genetic variation in Lycopersicon pimpinellifolium: evidence of evolutionary change in mating systems. Plant Systematics and Evolution, 127, 139–170.

Rosseel, Y. (2012) lavaan: An R Package for Structural Equation Modeling. Statistical Software, 48, 1–36.

Salathe, R. and Schmid-Hempel, P. (2011) The genotypic structure of a multi-host bumblebee parasite suggests a role for ecological niche overlap. PLoS One, 6, e22054.

Senthil-Kumar, M., Wang, K. and Mysore, K. (2013) At-CYP710A1 gene-mediated stigmasterol production plays a role in imparting temperature stress tolerance in Arabidopsis thaliana. Plant Signaling & Behavior, 8, e23142.

Sork, V., Davis, F., Westfall, R., Flint, A., Ikegami, M., Wang, H. and Grivet, D. (2010) Gene movement and genetic association with regional climate gradients in California valley oak (Quercus lobata Nee) in the face of climate change. Molecular Ecology, 19, 3806–3823.

Tanksley, S. (2004) The genetic, developmental, and molecular bases of fruit size and shape variation in tomato. The Plant cell, 16, 181–189.

Tanksley, S., Grandillo, S., Fulton, T., Zamir, D., Eshed, Y., Petiard, V., Lopez, J. and Beck-Bunn, T. (1996) Advanced backcross QTL analysis in a cross between an elite processing line of tomato and its wild relative L. pimpinellifolium. Theoretical and Applied Genetics, 92, 213–224.

Vucetich, J. and Walte, T. (2003) Spatial patterns of demography and genetic processes across the species’ range: Null hypotheses for landscape conservation genetics. Conservation Genetics, 4, 639–645.

Wang, I., Glor, R. and Losos, J. (2013) Quantifying the roles of ecology and geography in spatial genetic divergence. Ecology Letters, 16, 175–182.

Wiens, J. (1989) Spatial Scaling in Ecology. Functional Ecology, 3, 385–397.

van den Wollenberg, A. (1977) Redundancy analysis: an alternative for canonical correlation analysis. Psychometrika, 42, 207–219.

Wright, S. (1943) Isolation by distance. Genetics, 28, 114–138.

Yoder, J., Stanton-Geddes, J., Zhou, P., Briskine, R., Young, N. and Tiffin, P. (2012) Genomic signature of adaptation to climate in Medicago truncatula. Genetics, 196, 1263–1275.

Zhang, D., Du, Q., Zhang, Z., Jiao, X., Song, X. and Li, J. (2017) Vapour pressure deficit control in relation to water transport and water productivity in greenhouse tomato production during summer. Scientific Reports, 7, 43461.

Zomer, R., Trabucco, A., Bossio, D., van Straaten, O. and Verchot, L. (2008) Climate Change Mitigation: A Spatial Analysis of Global Land Suitability for Clean Development Mechanism Afforestation and Reforestation. Agric. Ecosystems and Envir, 126, 67–80.

Zuriaga, E., Blaanca, J., Cordero, L., Sifres, A., Blas-Cerdan, W., Morales, R. and Nuez, F. (2009) Genetic and bioclimatic variation in Solanum pimpinellifolium. Genetic Resources and Crop Evolution, 56, 39–51.

