## Supplemental info for "Regional differences in the abiotic environment contribute to genomic divergence within a wild tomato species"

Matthew J.S. Gibson

Leonie C. Moyle

**Table of Contents:**

| **Figure S1** | Page 2 |
| --- | --- |
| **Figure S2** | Page 3 |
| **Figure S3** | Page 4 |
| **Figure S4** | Page 5 |
| **Figure S5** | Page 6 |
| **Table S1** | Page 7 |
| **Table S2** | Page 8 |
| **Table S3** | Page 10 |
| **Table S4** | Page 11 |
| **Table S5** | Page 15 |
| **Table S6** | Page 17 |
| **Table S7** | Page 21 |
| **Table S8** | Page 21 |
| **Table S9** | Page 22 |
| **Table S10** | Page 22 |
| **Table S11** | Page 22 |
| **Table S12** | Page 22 |

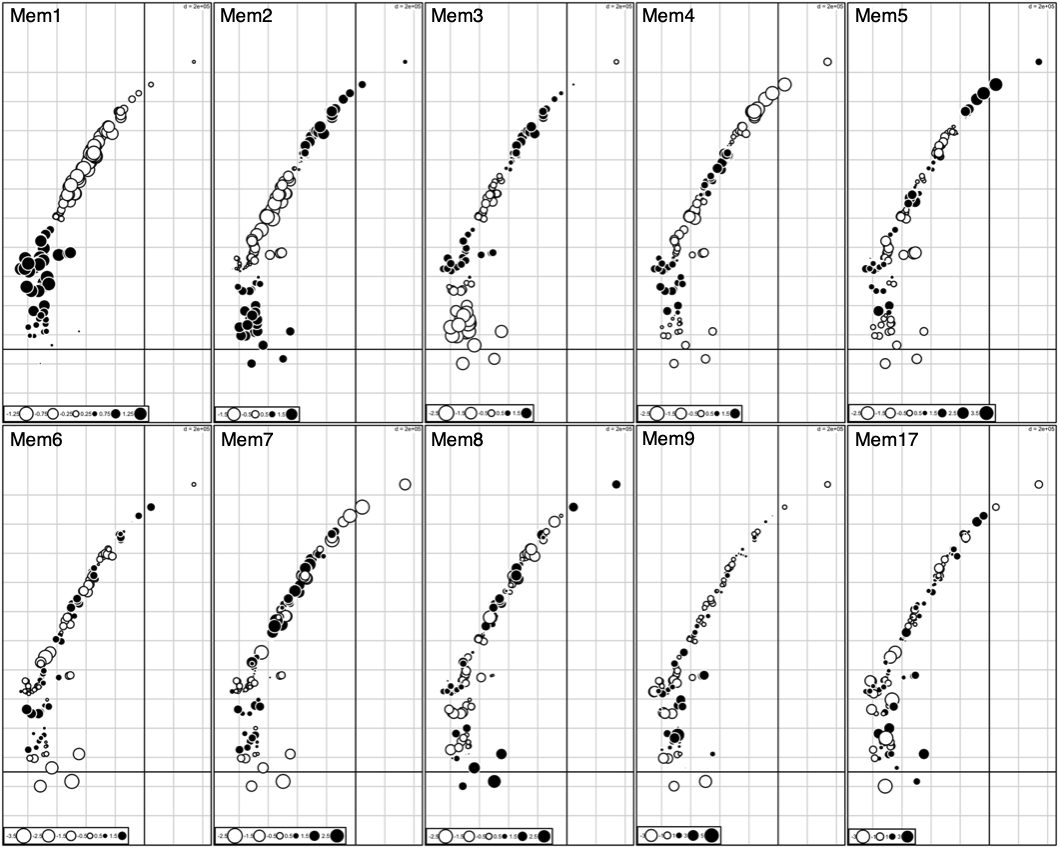

**Figure S1**: Bubble plots of the 10 dbMEM variables identified as significant. Variables 1-4 describe broad patterns of spatial structure and variables 5-10 describe fine-scale spatial variation. Color and size of the points correspond to the sign (+ or -) and magnitude of the dbMEM variables, respectively.

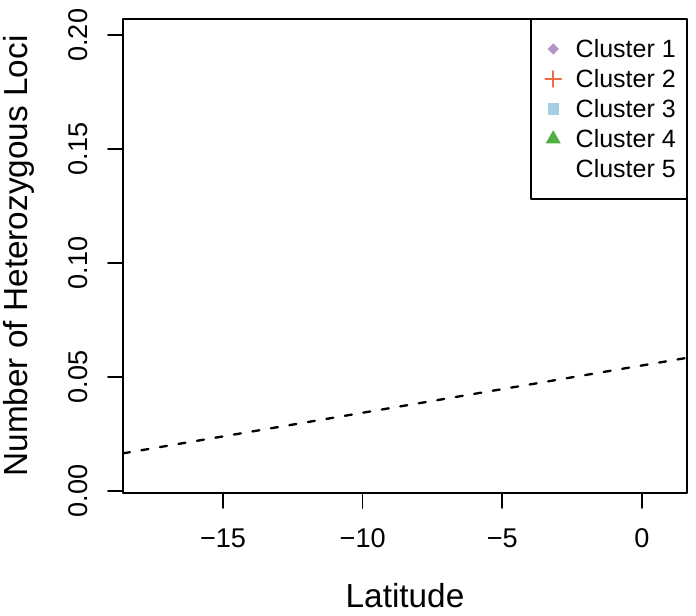

**Figure S2:** Plot of mean heterozygosity vs latitude. Heterozygosity increased towards the equator, concurrent with the center of the species range. Colors correspond to K-means clusters from PCA shown in Figure 2.

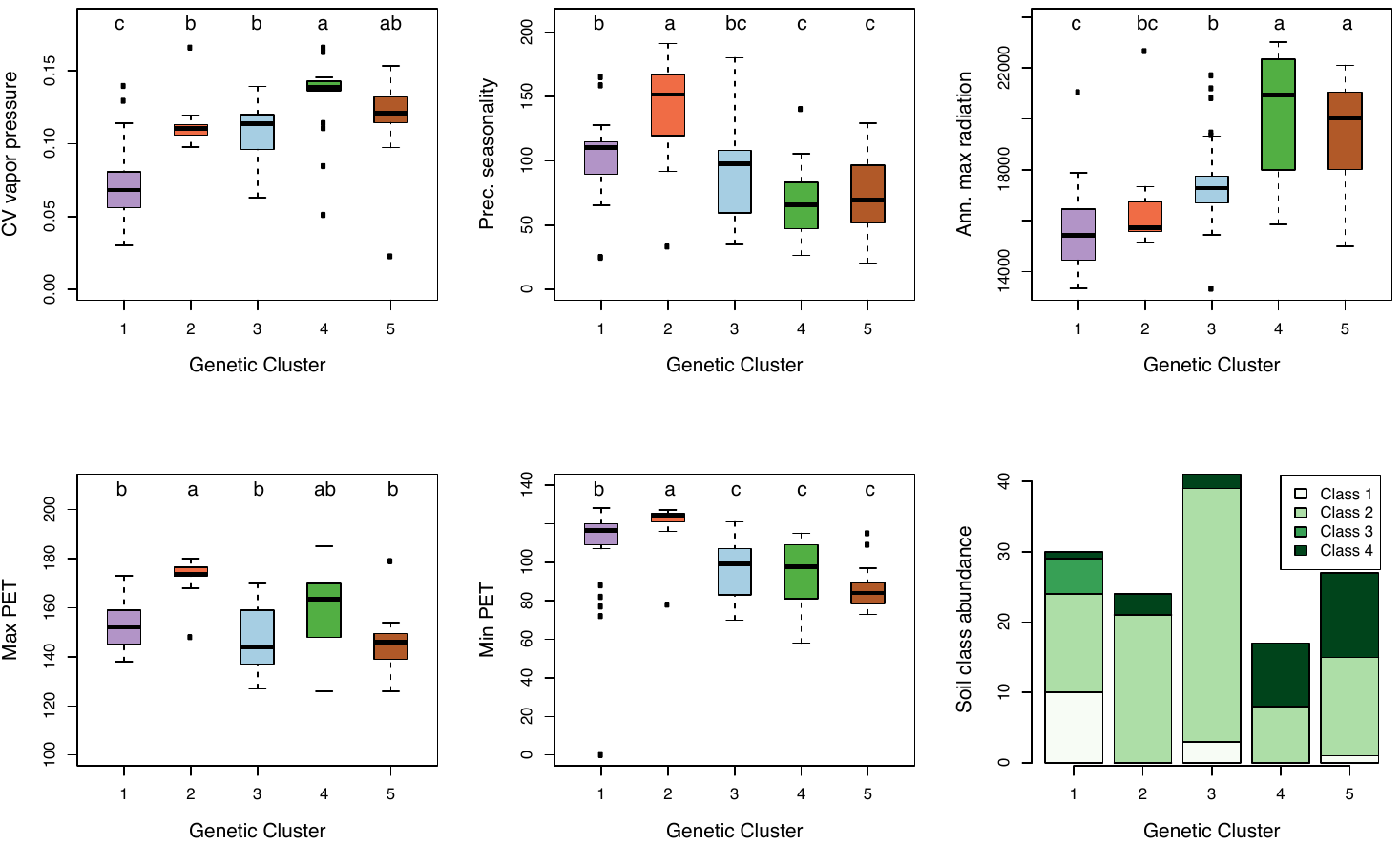

**Figure S3:** Climatic variation by regional genetic cluster. Colors correspond to K-means clusters from PCA shown in Figure 2. All panels were significant based on ANOVA (P < 1x10^-8^). Tukey HSD post-hoc comparison results are shown as letters above each box plot.

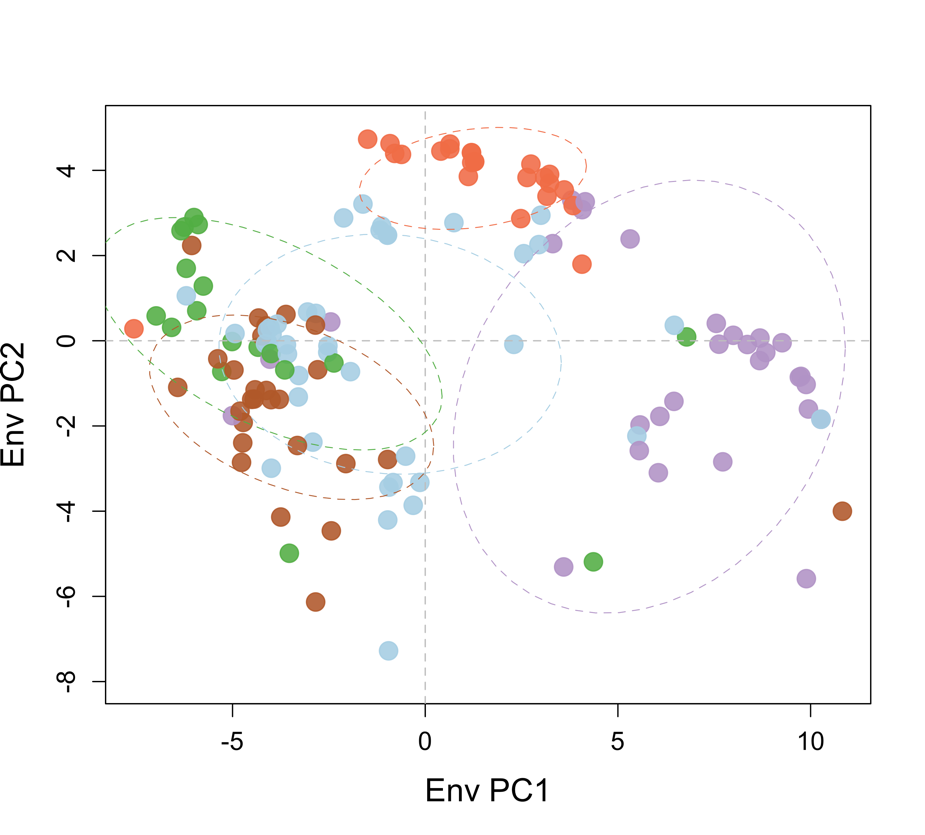

**Figure S4:** First two principal components of all 54 environmental variables. Colors correspond to K-means clusters shown in Figure 2.

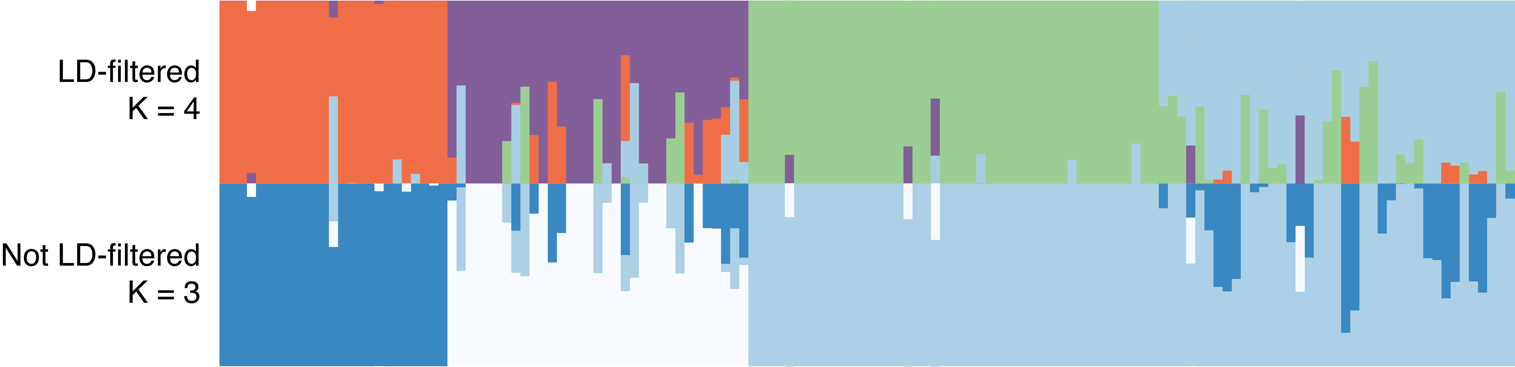

**Figure S5**: Comparison between fastStructure runs filtering (top) and not filtering (bottom) for LD, ordered from left to right by BIC cluster (Figure 2). The most likely K in the LD-filtered dataset was 4 whereas it was 3 is the unfiltered dataset. The major pattern on North-South structure is recapitulated in the unfiltered dataset, however original BIC clusters 3, 4, & 5 were largely merged into a single group (light blue; bottom panel). Note that different colors were used in the bottom panel to avoid confusion (white: BIC cluster 1; dark blue: BIC cluster 2; light blue: BIC cluster 3, 4, & 5).

**Table S1**: List of accessions used

[provided as separate spreadsheet]

**Table S2**: List of all environmental variables included in this study.

| **Variable** | **Database** |
| --- | --- |
| Annual mean temp | WorldClim |
| Mean diurnal range | WorldClim |
| Isothermality | WorldClim |
| Temperature seasonality | WorldClim |
| Max temp of warmest month | WorldClim |
| Min temp of warmest month | WorldClim |
| Temperature annual range | WorldClim |
| Mean temp of wettest quarter | WorldClim |
| Mean temp of driest quarter | WorldClim |
| Mean temp of warmest wuarter | WorldClim |
| Mean temp of coldest quarter | WorldClim |
| Annual precipitation | WorldClim |
| Precipitation of wettest month | WorldClim |
| Precipitation of driest month | WorldClim |
| Precipitation seasonality | WorldClim |
| Precipitation of wettest quarter | WorldClim |
| Precipitation of driest quarter | WorldClim |
| Precipitation of warmest quarter | WorldClim |
| Precipitation of coldest quarter | WorldClim |
| Annual average radiation | WorldClim |
| Annual maximum radiation | WorldClim |
| Annual minimum radiation | WorldClim |
| Annual variation in solar radiation | WorldClim |
| Annual average vapor pressure | WorldClim |
| Annual max vapor pressure | WorldClim |
| Annual minimum vapor pressure | WorldClim |
| Annual variation in vapor pressure | WorldClim |
| Annual average wind speed | WorldClim |
| Annual max wind speed | WorldClim |
| Annual minimum wind speed | WorldClim |
| Annual variation in wind speed | WorldClim |
| Annual heat:moisture index | Climate South America (CSA) |
| Degree days above 5 C | Climate South America (CSA) |
| Degree days below 18 C | Climate South America (CSA) |
| Degree days above 18 C | Climate South America (CSA) |
| Number of frost free days | Climate South America (CSA) |
| Extreme minimum temperature over 30 years | Climate South America (CSA) |
| Hargreaves reference evaporation | Climate South America (CSA) |
| Hargreaves climatic moisture deficit | Climate South America (CSA) |
| Annual aridity | CGIAR |
| Annual potential evapotranspiration | CGIAR |
| Annual max potential evapotranspiration | CGIAR |
| Annual min potential evapotranspiration | CGIAR |
| Annual variation potential evapotranspiration | CGIAR |
| Annual Actual evapotranspiration | CGIAR |
| Annual soil water stress. Mean of months | CGIAR |
| Annual max soil water stress | CGIAR |
| Annual min soil water stress | CGIAR |
| Annual variation soil water stress | CGIAR |
| Priestley-Taylor Alpha Coefficient | CGIAR |
| Bulk soil density (fine earth fraction) | SoilGrids |
| Cation exchange capacity | SoilGrids |
| pH | SoilGrids |
| Soil Texture | SoilGrids |

**Table S3**: Stacks ref_map assembly summary

| **Statistic** | **Value** |
| --- | --- |
| Bam records | 444,288,642 |
| Primary alignments retained | 431,111,093 |
| Records per sample | 2,776,804 (17,586-13,836,425) |
| Assembled/genotyped loci | 458,945 |
| Effective per-sample coverage | 66.1x (sd = 36.7x) |
| Mean sites per locus | 139.5 |

**Table S4**: Stacks per-sample assembly summary

| **Sample** | **Bam records** | **% kept** | **# Loci** | **Mean Coverage** |
| --- | --- | --- | --- | --- |
| 0121-101 | 860527 | 0.971 | 33899 | 24.648 |
| 0122-101 | 1630820 | 0.969 | 49273 | 32.075 |
| 0373-102 | 3329913 | 0.97 | 45773 | 70.533 |
| 0375-101 | 2126539 | 0.963 | 64386 | 31.809 |
| 0418-101 | 2143602 | 0.975 | 62968 | 33.189 |
| 0420-101 | 1719200 | 0.976 | 58616 | 28.615 |
| 0442-101 | 4607456 | 0.972 | 88923 | 50.381 |
| 0443-101 | 2497478 | 0.969 | 69306 | 34.922 |
| 0753-101 | 1107015 | 0.967 | 48569 | 22.039 |
| 0772-101 | 2065633 | 0.97 | 53973 | 37.118 |
| 1248-101 | 2745125 | 0.978 | 60529 | 44.377 |
| 1256-101 | 3632845 | 0.975 | 69302 | 51.117 |
| 1280-101 | 3120897 | 0.974 | 59962 | 50.697 |
| 1301-101 | 2155029 | 0.975 | 61959 | 33.905 |
| 1381-101 | 2114032 | 0.97 | 68611 | 29.901 |
| 1416-101 | 313218 | 0.978 | 27478 | 11.143 |
| 1428-101 | 2700077 | 0.979 | 69663 | 37.951 |
| 1469-101 | 1457202 | 0.971 | 58027 | 24.386 |
| 1470-101 | 3475525 | 0.972 | 69800 | 48.403 |
| 1472-101 | 12103090 | 0.969 | 102173 | 114.79 |
| 1478-101 | 989959 | 0.975 | 38844 | 24.838 |
| 1520-101 | 4548266 | 0.974 | 59564 | 74.396 |
| 1547-101 | 874210 | 0.977 | 45564 | 18.748 |
| 1561-101 | 2172059 | 0.975 | 53624 | 39.495 |
| 1562-101 | 2288002 | 0.972 | 53428 | 41.645 |
| 1572-101 | 4330309 | 0.975 | 70438 | 59.918 |
| 1576-101 | 4227007 | 0.965 | 88818 | 45.908 |
| 1579-101 | 2739722 | 0.972 | 58816 | 45.268 |
| 1584-101 | 3938382 | 0.97 | 74897 | 51.014 |
| 1586-101 | 3544559 | 0.968 | 72654 | 47.244 |
| 1589-12 | 823915 | 0.969 | 35416 | 22.532 |
| 1598-101 | 4697846 | 0.975 | 64759 | 70.721 |
| 1600-101 | 2096174 | 0.969 | 61406 | 33.065 |
| 1601-101 | 301039 | 0.961 | 23332 | 12.399 |
| 1602-101 | 2736356 | 0.969 | 68369 | 38.786 |
| 1603-101 | 5257707 | 0.974 | 73130 | 69.994 |
| 1604-101 | 2492691 | 0.943 | 57000 | 41.24 |
| 1605-101 | 3183592 | 0.977 | 72458 | 42.94 |
| 1611-101 | 2652137 | 0.97 | 60598 | 42.432 |
| 1614-101 | 2035135 | 0.974 | 52730 | 37.608 |
| 1615-101 | 2245827 | 0.976 | 57469 | 38.127 |
| 1635-101 | 5004334 | 0.965 | 87139 | 55.392 |
| 1636-101 | 1900561 | 0.972 | 60673 | 30.458 |
| 1637-101 | 1664781 | 0.974 | 51442 | 31.533 |
| 1651-101 | 266205 | 0.976 | 29007 | 8.959 |
| 1659-101 | 5447399 | 0.975 | 79921 | 66.446 |
| 1660-101 | 1923801 | 0.965 | 54283 | 34.193 |
| 1661-101 | 2289988 | 0.975 | 61667 | 36.212 |
| 1670-101 | 3133778 | 0.975 | 61588 | 49.586 |
| 1683-101 | 3318632 | 0.966 | 69450 | 46.168 |
| 1684-101 | 3191539 | 0.973 | 44759 | 69.411 |
| 1685-101 | 2731893 | 0.973 | 62235 | 42.725 |
| 1686-101 | 3440511 | 0.974 | 69004 | 48.576 |
| 1687-101 | 4310773 | 0.975 | 73979 | 56.835 |
| 1688-101 | 4800347 | 0.974 | 78301 | 59.725 |
| 1689-101 | 1127857 | 0.976 | 42640 | 25.82 |
| 1690-101 | 2555961 | 0.973 | 61518 | 40.42 |
| 1697-101 | 5430669 | 0.975 | 80374 | 65.871 |
| 1728-101 | 5326058 | 0.959 | 48284 | 105.82 |
| 1729-101 | 5544196 | 0.976 | 84469 | 64.066 |
| 1921-101 | 2499661 | 0.968 | 62252 | 38.871 |
| 1923-101 | 3376990 | 0.972 | 65166 | 50.35 |
| 1924-101 | 3200102 | 0.974 | 66391 | 46.935 |
| 1933-101 | 2357305 | 0.974 | 64160 | 35.795 |
| 1936-101 | 13836425 | 0.973 | 80387 | 167.542 |
| 1950-101 | 4561743 | 0.971 | 70862 | 62.512 |
| 1987-101 | 290857 | 0.964 | 26270 | 10.678 |
| 2069-101 | 3869190 | 0.975 | 71933 | 52.432 |
| 2093-101 | 2700416 | 0.976 | 72879 | 36.181 |
| 2102-101 | 2219077 | 0.971 | 69103 | 31.176 |
| 2181-101 | 4189761 | 0.971 | 76704 | 53.044 |
| 2186-101 | 2449823 | 0.976 | 64750 | 36.929 |
| 2401-101 | 2238459 | 0.968 | 67265 | 32.229 |
| 2412-101 | 555326 | 0.955 | 33384 | 15.883 |
| 251318-101 | 2181450 | 0.974 | 57605 | 36.898 |
| 251319-101 | 977057 | 0.975 | 45405 | 20.971 |
| 251320-101 | 3661717 | 0.971 | 78245 | 45.449 |
| 251321-101 | 2735553 | 0.974 | 55029 | 48.435 |
| 2544-101 | 4029122 | 0.974 | 70886 | 55.365 |
| 2576-101 | 2840786 | 0.968 | 64062 | 42.918 |
| 2578-101 | 1676259 | 0.971 | 55032 | 29.572 |
| 2649-101 | 420490 | 0.962 | 29144 | 13.873 |
| 2650-101 | 229441 | 0.978 | 19064 | 11.776 |
| 2725-101 | 4544274 | 0.975 | 74845 | 59.176 |
| 2832-101 | 3536608 | 0.97 | 76226 | 44.996 |
| 2839-101 | 4515964 | 0.966 | 75744 | 57.575 |
| 2852-101 | 1051515 | 0.97 | 44880 | 22.728 |
| 2854-101 | 1761127 | 0.977 | 52639 | 32.686 |
| 2857-101 | 2620451 | 0.978 | 73055 | 35.095 |
| 2915-101 | 3738875 | 0.972 | 68961 | 52.7 |
| 2933-101 | 73476 | 0.951 | 12699 | 5.504 |
| 2983-101 | 4009330 | 0.974 | 69299 | 56.376 |
| 3123-101 | 1426303 | 0.968 | 57828 | 23.874 |
| 365909-101 | 1356728 | 0.977 | 54425 | 24.349 |
| 365910-101 | 3475812 | 0.979 | 68715 | 49.517 |
| 365911-101 | 17586 | 0.935 | 6449 | 2.551 |
| 365912-101 | 2690411 | 0.978 | 56733 | 46.383 |
| 365914-101 | 2912501 | 0.975 | 77470 | 36.637 |
| 365915-101 | 2507336 | 0.976 | 71185 | 34.393 |
| 365916-101 | 1756143 | 0.975 | 61231 | 27.969 |
| 365917-101 | 1106543 | 0.977 | 43360 | 24.93 |
| 365918-101 | 3367378 | 0.976 | 83068 | 39.566 |
| 365958-101 | 4835275 | 0.972 | 72102 | 65.217 |
| 365960-101 | 2534529 | 0.974 | 60177 | 41.017 |
| 365961-101 | 2291525 | 0.975 | 56735 | 39.39 |
| 365962-101 | 3773949 | 0.971 | 71611 | 51.174 |
| 365964-101 | 3201558 | 0.973 | 71511 | 43.553 |
| 365965-101 | 3722221 | 0.975 | 55326 | 65.626 |
| 365966-101 | 1822739 | 0.976 | 49693 | 35.796 |
| 379021-101 | 1778550 | 0.974 | 55735 | 31.077 |
| 379023-101 | 2888964 | 0.973 | 66555 | 42.252 |
| 379024-101 | 1718608 | 0.97 | 50560 | 32.978 |
| 379025-101 | 4022951 | 0.975 | 66419 | 59.073 |
| 379026-101 | 4340700 | 0.977 | 74945 | 56.579 |
| 379059-101 | 2421589 | 0.978 | 61995 | 38.185 |
| 390519-101 | 1666679 | 0.973 | 58615 | 27.678 |
| 390688-101 | 1752668 | 0.974 | 60648 | 28.143 |
| 390689-101 | 4512953 | 0.973 | 74530 | 58.917 |
| 390690-101 | 2646066 | 0.975 | 21929 | 117.596 |
| 390691-101 | 3381952 | 0.973 | 70517 | 46.682 |
| 390695-101 | 423532 | 0.973 | 28031 | 14.7 |
| 390696-101 | 3692047 | 0.972 | 70611 | 50.849 |
| 390699-101 | 2130632 | 0.972 | 62135 | 33.314 |
| 390700-101 | 3748237 | 0.972 | 66812 | 54.525 |
| 390701-101 | 2286853 | 0.972 | 69124 | 32.141 |
| 390705-101 | 2541073 | 0.973 | 74752 | 33.063 |
| 390706-101 | 5667961 | 0.975 | 82057 | 67.372 |
| 390707-101 | 2150065 | 0.973 | 62573 | 33.427 |
| 390716-101 | 649115 | 0.974 | 45899 | 13.768 |
| 390718-101 | 2263338 | 0.973 | 67281 | 32.74 |
| 390721-101 | 1990569 | 0.974 | 62742 | 30.907 |
| 390722-101 | 2426132 | 0.975 | 60309 | 39.233 |
| 390728-101 | 4222460 | 0.975 | 71596 | 57.501 |
| 390748-102 | 1619440 | 0.974 | 56297 | 28.025 |
| 407533-101 | 1313574 | 0.97 | 59705 | 21.342 |
| 407534-101 | 3650109 | 0.974 | 71089 | 50.01 |
| 407535-103 | 2199289 | 0.973 | 66777 | 32.044 |
| 407537-101 | 3609456 | 0.973 | 62715 | 55.979 |
| 407538-101 | 2500804 | 0.969 | 64978 | 37.293 |
| 407544-101 | 1986374 | 0.969 | 61346 | 31.374 |
| 407550-101 | 4712208 | 0.976 | 70940 | 64.811 |
| 407551-101 | 2952304 | 0.966 | 65073 | 43.827 |
| 407552-101 | 2026851 | 0.975 | 50963 | 38.76 |
| 407553-101 | 3324274 | 0.975 | 64736 | 50.057 |
| 407554-101 | 1600623 | 0.971 | 59342 | 26.196 |
| 503517-101 | 41587 | 0.973 | 12061 | 3.356 |
| 503519-101 | 2150172 | 0.973 | 60312 | 34.683 |
| 503521-101 | 6484748 | 0.975 | 88499 | 71.477 |
| 503522-101 | 2816838 | 0.971 | 71276 | 38.385 |
| 503523-101 | 2419634 | 0.97 | 61510 | 38.151 |
| 503524-101 | 1960823 | 0.979 | 61581 | 31.178 |

**Table S5**: Summary of genotype filters

| **Filter #** | **Filter applied** | **Script calls** | **# of SNPs** | **Used for** |
| --- | --- | --- | --- | --- |
| Initial Stacks calls | - | Stacks ref_map, populations | 366,466 | - |
| 1 | Remove sites mapped to unassembled contigs | grep -Ev 'SL3.0ch00' populations.snps.vcf | 359,796 | - |
| 2 | Change calls made with < 4 reads to missing | vcftools --vcf populations.snps.filter1.vcf --recode --recode-INFO-all --out populations.snps.filter2 --minDP 4 | 359,796 | - |
| 3 | Remove sites genotyped in fewer than 30% of individuals | vcftools --vcf populations.snps.filter2.pimp.vcf --recode --recode-INFO-all --out populations.snps.filter3.pimp --max-missing-count 120 | 44,533 | - |
| 4 | Remove sites with heterozygosity > 0.6 | Tassel GUI filter: Max heterozygous = 0.6 | 44,243 | - |
| 5 | Remove individuals with > 60% missing data | vcftools --vcf populations.snps.filter4.pimp.vcf --remove-indv 365911-101 --remove-indv 1601-101 --remove-indv 1416-101 --remove-indv 1987-101 --remove-indv 2649-101 --remove-indv 2933-101 --remove-indv 390695-101 --remove-indv 503517-101 --recode --recode-INFO-all --out populations.snps.filter5.pimp | 44,064 | RDA genome-environment scan |
| 6 | Remove SNPs in high LD | bcftools +prune -l 0.9 -w 1000 populations.snps.filter5.pimp.recode.vcf -Ov -o populations.snps.filter6.pimp.vcf | 17,358 | - |
| 7 | Remove sites genotyped in less than 95% of individuals | vcftools --vcf populations.snps.filter6.pimp.vcf --recode --recode-INFO-all --out populations.snps.filter7.pimp --max-missing-count 8 | 7,317 | PCA, fastStructure |
| 7 | Remove SNPs with any missing data | vcftools --vcf populations.snps.filter7.pimp.vcf --recode --recode-INFO-all --out populations.snps.filter8.pimp --max-missing-count 0 | 6,830 | RDA variance partitioning, SEM, GDM |

**Table S6**: Full summary of SEM model fit

|  | **Term** | **Est** | **SE** | **z** | **CI lower** | **CI upper** |
| --- | --- | --- | --- | --- | --- | --- |
| **Latent** | bio_1 | 1 | 0 | NA | 1 | 1 |
|  | bio_2 | 0.292 | 0.030 | 9.514 | 0.231 | 0.352 |
|  | bio_3 | 1.724 | 0.063 | 26.995 | 1.599 | 1.849 |
|  | bio_4 | 1.869 | 0.067 | 27.815 | 1.736 | 2.000 |
|  | bio_5 | 0.361 | 0.024 | 14.606 | 0.312 | 0.409 |
|  | bio_6 | 1.568 | 0.035 | 43.599 | 1.497 | 1.633 |
|  | bio_7 | 1.168 | 0.046 | 25.170 | 1.077 | 1.259 |
|  | bio_8 | 0.315 | 0.026 | 11.745 | 0.262 | 0.367 |
|  | bio_9 | 0.749 | 0.020 | 37.148 | 0.709 | 0.788 |
|  | bio_10 | 0.515 | 0.019 | 25.899 | 0.475 | 0.553 |
|  | bio_11 | 1.682 | 0.038 | 43.795 | 1.606 | 1.756 |
|  | bio_12 | 2.766 | 0.102 | 26.91 | 2.564 | 2.967 |
|  | bio_13 | 2.786 | 0.103 | 26.989 | 2.584 | 2.988 |
|  | bio_14 | 1.53 | 0.058 | 26.35 | 1.416 | 1.644 |
|  | bio_15 | 0.671 | 0.035 | 18.758 | 0.600 | 0.741 |
|  | bio_16 | 2.812 | 0.105 | 26.761 | 2.606 | 3.019 |
|  | bio_17 | 1.520 | 0.056 | 26.749 | 1.408 | 1.631 |
|  | bio_18 | 2.594 | 0.094 | 27.515 | 2.409 | 2.778 |
|  | bio_19 | 1.070 | 0.049 | 21.638 | 0.973 | 1.166 |
|  | ann_avg_rad | 1.404 | 0.053 | 26.016 | 1.298 | 1.509 |
|  | ann_max_rad | 1.481 | 0.055 | 26.829 | 1.373 | 1.589 |
|  | ann_min_rad | 0.260 | 0.030 | 8.579 | 0.201 | 0.320 |
|  | cv_rad | 0.325 | 0.031 | 10.401 | 0.263 | 0.386 |
|  | ann_avg_vapr | 1.483 | 0.037 | 39.459 | 1.409 | 1.556 |
|  | ann_max_vapr | 1.129 | 0.022 | 50.454 | 1.085 | 1.173 |
|  | ann_min_vapr | 1.529 | 0.038 | 39.324 | 1.453 | 1.606 |
|  | cv_vapr | 2.249 | 0.079 | 28.364 | 2.094 | 2.405 |
|  | ann_avg_wind | 1.601 | 0.070 | 22.745 | 1.463 | 1.739 |
|  | ann_max_wind | 1.601 | 0.070 | 22.822 | 1.464 | 1.739 |
|  | ann_min_wind | 1.399 | 0.06 | 21.452 | 1.271 | 1.527 |
|  | cv_wind | 0.563 | 0.034 | 16.475 | 0.496 | 0.630 |
|  | AHM | 2.799 | 0.102 | 27.217 | 2.597 | 3.000 |
|  | DD5 | 1.171 | 0.023 | 48.988 | 1.124 | 1.218 |
|  | DD_18 | 0.616 | 0.019 | 32.072 | 0.578 | 0.653 |
|  | DD18 | 1.719 | 0.055 | 31.158 | 1.611 | 1.827 |
|  | NFFD | 0.212 | 0.032 | 6.638 | 0.149 | 0.275 |
|  | EMT | 0.530 | 0.020 | 25.460 | 0.489 | 0.570 |
|  | Eref | 0.332 | 0.027 | 12.013 | 0.278 | 0.387 |
|  | CMD | 1.696 | 0.060 | 28.077 | 1.577 | 1.814 |
|  | ann_ai | 2.878 | 0.105 | 27.164 | 2.670 | 3.086 |
|  | ann_pet | 0.477 | 0.028 | 16.783 | 0.421 | 0.533 |
|  | max_pet | 0.195 | 0.026 | 7.280 | 0.142 | 0.248 |
|  | min_pet | 0.854 | 0.035 | 23.922 | 0.784 | 0.924 |
|  | cv_pet | 2.171 | 0.073 | 29.465 | 2.026 | 2.315 |
|  | ann_aet | 2.878 | 0.105 | 27.340 | 2.671 | 3.084 |
|  | ann_swt | 1.933 | 0.075 | 25.571 | 1.784 | 2.081 |
|  | max_swt | 0.663 | 0.036 | 18.073 | 0.591 | 0.734 |
|  | min_swt | 1.713 | 0.061 | 27.659 | 1.592 | 1.834 |
|  | cv_swt | 1.133 | 0.051 | 22.061 | 1.032 | 1.233 |
|  | ptac | 2.594 | 0.095 | 27.273 | 2.408 | 2.781 |
|  | bdfe | 1.537 | 0.053 | 28.761 | 1.432 | 1.642 |
|  | cec | 0.617 | 0.037 | 16.248 | 0.542 | 0.691 |
|  | swc | 0.484 | 0.032 | 14.767 | 0.420 | 0.548 |
|  | ph | 2.164 | 0.080 | 26.896 | 2.006 | 2.321 |
|  | txt.4 | 1.089 | 0.050 | 21.534 | 0.990 | 1.188 |
|  | txt.6 | 0.564 | 0.036 | 15.504 | 0.492 | 0.635 |
|  | txt.7 | 0.642 | 0.039 | 16.441 | 0.566 | 0.719 |
|  | txt.9 | 0.468 | 0.036 | 12.988 | 0.398 | 0.539 |
|  | geo.dist | 1 | 0 | NA | 1 | 1 |
| **Regressions** | gen.dist~env | 0.788 | 0.043 | 18.066 | 0.702 | 0.873 |
|  | gen.dist~geo | 0.17 | 0.011 | 14.523 | 0.150 | 0.197 |
| **Covariances** | env~geo | 0.219 | 0.010 | 21.967 | 0.200 | 0.239 |
| **Variances** | geo | 0.219 | 0.010 | 21.967 | 0.200 | 0.239 |
|  | env | 0.112 | 0.008 | 13.967 | 0.096 | 0.128 |
|  | geo | 0.999 | 0.013 | 72.773 | 0.972 | 1.026 |
|  | bio_1 | 0.887 | 0.042 | 20.956 | 0.804 | 0.970 |
|  | bio_2 | 0.990 | 0.013 | 75.511 | 0.964 | 1.015 |
|  | bio_3 | 0.664 | 0.010 | 60.748 | 0.643 | 0.685 |
|  | bio_4 | 0.606 | 0.008 | 67.590 | 0.588 | 0.623 |
|  | bio_5 | 0.985 | 0.031 | 30.838 | 0.922 | 1.047 |
|  | bio_6 | 0.722 | 0.021 | 34.341 | 0.681 | 0.763 |
|  | bio_7 | 0.845 | 0.014 | 59.574 | 0.818 | 0.873 |
|  | bio_8 | 0.988 | 0.018 | 52.292 | 0.951 | 1.025 |
|  | bio_9 | 0.936 | 0.026 | 35.641 | 0.885 | 0.988 |
|  | bio_10 | 0.969 | 0.033 | 29.191 | 0.904 | 1.035 |
|  | bio_11 | 0.680 | 0.020 | 33.767 | 0.641 | 0.720 |
|  | bio_12 | 0.137 | 0.003 | 40.948 | 0.130 | 0.143 |
|  | bio_13 | 0.124 | 0.003 | 40.062 | 0.118 | 0.130 |
|  | bio_14 | 0.735 | 0.014 | 51.722 | 0.707 | 0.763 |
|  | bio_15 | 0.949 | 0.013 | 69.279 | 0.922 | 0.975 |
|  | bio_16 | 0.107 | 0.003 | 32.742 | 0.101 | 0.113 |
|  | bio_17 | 0.739 | 0.010 | 67.260 | 0.717 | 0.760 |
|  | bio_18 | 0.240 | 0.005 | 43.859 | 0.230 | 0.251 |
|  | bio_19 | 0.870 | 0.011 | 74.274 | 0.847 | 0.893 |
|  | ann_avg_rad | 0.777 | 0.010 | 71.800 | 0.756 | 0.798 |
|  | ann_max_rad | 0.752 | 0.009 | 77.481 | 0.733 | 0.771 |
|  | ann_min_rad | 0.992 | 0.015 | 63.339 | 0.961 | 1.022 |
|  | cv_rad | 0.987 | 0.016 | 61.618 | 0.956 | 1.019 |
|  | ann_avg_vapr | 0.751 | 0.018 | 40.139 | 0.715 | 0.788 |
|  | ann_max_vapr | 0.855 | 0.020 | 37.367 | 0.811 | 0.900 |
|  | ann_min_vapr | 0.736 | 0.019 | 37.210 | 0.697 | 0.774 |
|  | cv_vapr | 0.429 | 0.006 | 63.030 | 0.415 | 0.442 |
|  | ann_avg_wind | 0.710 | 0.010 | 68.549 | 0.690 | 0.731 |
|  | ann_max_wind | 0.710 | 0.010 | 66.884 | 0.689 | 0.731 |
|  | ann_min_wind | 0.779 | 0.010 | 73.168 | 0.758 | 0.799 |
|  | cv_wind | 0.964 | 0.015 | 62.054 | 0.933 | 0.994 |
|  | AHM | 0.116 | 0.003 | 32.730 | 0.109 | 0.123 |
|  | DD5 | 0.845 | 0.016 | 49.901 | 0.811 | 0.878 |
|  | DD_18 | 0.957 | 0.016 | 56.487 | 0.923 | 0.990 |
|  | DD18 | 0.666 | 0.010 | 65.954 | 0.646 | 0.686 |
|  | NFFD | 0.994 | 0.058 | 16.986 | 0.880 | 1.109 |
|  | EMT | 0.968 | 0.020 | 48.004 | 0.928 | 1.007 |
|  | Eref | 0.987 | 0.015 | 65.433 | 0.957 | 1.016 |
|  | CMD | 0.675 | 0.014 | 47.645 | 0.647 | 0.703 |
|  | ann_ai | 0.065 | 0.003 | 16.743 | 0.057 | 0.073 |
|  | ann_pet | 0.974 | 0.014 | 65.480 | 0.945 | 1.003 |
|  | max_pet | 0.995 | 0.050 | 19.586 | 0.895 | 1.095 |
|  | min_pet | 0.917 | 0.027 | 32.907 | 0.862 | 0.972 |
|  | cv_pet | 0.468 | 0.007 | 65.070 | 0.454 | 0.482 |
|  | ann_aet | 0.065 | 0.003 | 18.125 | 0.058 | 0.072 |
|  | ann_swt | 0.570 | 0.009 | 57.975 | 0.558 | 0.598 |
|  | max_swt | 0.950 | 0.012 | 76.405 | 0.925 | 0.974 |
|  | min_swt | 0.668 | 0.007 | 89.816 | 0.654 | 0.683 |
|  | cv_swt | 0.855 | 0.011 | 72.130 | 0.831 | 0.878 |
|  | ptac | 0.240 | 0.004 | 49.526 | 0.231 | 0.250 |
|  | bdfe | 0.733 | 0.015 | 48.467 | 0.703 | 0.762 |
|  | cec | 0.956 | 0.017 | 54.409 | 0.922 | 0.991 |
|  | swc | 0.97 | 0.017 | 56.028 | 0.939 | 1.007 |
|  | ph | 0.471 | 0.007 | 65.457 | 0.457 | 0.485 |
|  | txt.4 | 0.86 | 0.014 | 58.721 | 0.837 | 0.895 |
|  | txt.6 | 0.964 | 0.004 | 225.240 | 0.955 | 0.972 |
|  | txt.7 | 0.953 | 0.031 | 30.108 | 0.891 | 1.015 |
|  | txt.9 | 0.975 | 0.008 | 115.088 | 0.958 | 0.991 |
|  | geo.dist | 0 | 0 | NA | 0 | 0 |
|  | gen.dist | 0.839 | 0.013 | 60.734 | 0.812 | 0.866 |

**Table S7:** Full summary of GDM models

| **Model** | **Null deviance** | **GDM deviance** | **Percent deviance explained** | **Intercept** |
| --- | --- | --- | --- | --- |
| Environment | 863.28 | 611.06 | 29.21 | 0.96 |
| Space | 863.28 | 657.36 | 23.85 | 0.95 |
| Environment+Space | 863.28 | 602.45 | 30.21 | 0.90 |

**Table S8:** Variable contributions for environment GDM model. 3-spline fit.

| **Predictor** | **Coefficient 1** | **Coefficient 2** | **Coefficient 3** |
| --- | --- | --- | --- |
| DD18 | 0 | 0.692 | 0 |
| Bio7 | 0 | 0 | 0.065 |
| ann_max_rad | 0 | 0.108 | 0.130 |
| max_swt | 0 | 0.019 | 0.014 |
| cv_eind | 0.065 | 0 | 0 |
| Bio4 | 0 | 0 | 0 |
| ann_avg_wind | 0 | 0 | 0.112 |
| ann_max_wind | 0 | 0 | 0 |
| max_pet | 0 | 0 | 0.067 |
| CMD | 0 | 0 | 0 |
| cv_vapr | 0 | 0.212 | 0 |
| EMT | 0 | 0 | 0 |
| Bio3 | 0.022 | 0 | 0 |
| Bio2 | 0 | 0.009 | 0.003 |

**Table S9**: Variable contributions for environment + space GDM model. 3-spline fit.

| **Predictor** | **Coefficient1** | **Coefficient2** | **Coefficient3** |
| --- | --- | --- | --- |
| Geographic | 0.205 | 0.168 | 0.083 |
| DD18 | 0 | 0.566 | 0 |
| Bio7 | 0 | 0 | 0.041 |
| Ann_max_rad | 0 | 0.054 | 0.057 |
| max_swt | 0 | 0 | 0.028 |
| cv_eind | 0.014 | 0 | 0 |
| Bio4 | 0 | 0.033 | 0 |
| ann_avg_wind | 0 | 0 | 0.092 |
| ann_max_wind | 0 | 0 | 0 |
| max_pet | 0 | 0 | 0.038 |
| CMD | 0 | 0 | 0 |
| cv_vapr | 0 | 0.027 | 0 |
| EMT | 0 | 0 | 0 |
| Bio3 | 0 | 0.024 | 0 |
| Bio2 | 0 | 0 | 0.005 |

**Table S10:** Correlations among PCs, dbMEM variables, and environmental variables. Correlation coefficients are shown in the lower diagonal. Corrected P-values are shown in the upper diagonal.

[provided as separate spreadsheet]

**Table S11:** Full list of outlier SNPs and their annotations.

[provided as separate spreadsheet]

**Table S12:** Collated environmental data for each accession. See **Table S2** for descriptions of each variable and their source database.

[provided as separate spreadsheet]
